# Functional organization of the neonatal thalamus across development depicted by functional MRI

**DOI:** 10.1101/2025.01.18.633703

**Authors:** Hamza Kebiri, Farnaz Delavari, Dimitri Van De Ville, João Jorge, Meritxell Bach Cuadra

## Abstract

The thalamus is a central component of the brain that is involved in a variety of functions, from sensory processing to high-order cognition. Its structure and function in the first weeks of extrauterine life, including its connections to different cortical and subcortical areas, have not yet been widely explored. Here, we used resting state functional magnetic resonance imaging data of 730 newborns from the developing Human Connectome Project to study the functional organization of the thalamus from 37 to 44 post-conceptual weeks. We introduce KNIT: K-means for Nuclei in Infant Thalamus. The framework employs a highly granular vector space of 40 features, each corresponding to functional connectivity to a brain region, using *k-means* clustering and uncertainty quantification through bootstrapping to delineate thalamic units. Although the different clusters showed common patterns of increased connectivity to the superior temporal gyrus, the parietal, and the frontal cortex, implying an expected decrease in specialization at that age, they also show some specificity. That is, a pulvinar unit was identified, similar to the adult thalamus. Ventrolateral motor and medial salience units were also highlighted. The latter appeared around 41 weeks of age, while the former showed at least from 37 weeks, but had a decrease in volume through age, replaced mostly by a dominant dorsal thalamic unit. We also observed an increase in clustering robustness and in hemispheric bilateral symmetry with age, suggesting more specialized functional units. We also found a burst in global thalamic connectivity around 41 weeks. Finally, we demonstrate the benefits of this method in terms of granularity compared to the more conventional winner-takes-all approach.

## 1 Introduction

The thalamus is a prominent structure of the brain involved in both low and high-order brain functions. It is often referred to as a relay center of the brain, transmitting bottom-up signals from all sensory modalities (except olfaction), to respective cortical areas (Hwang et al., 2017) for dedicated processing. Additionally, the thalamus serves as a hub integrating cortical and subcortical information. This makes it intricately involved not only in sensory processing and vital cardio-respiratory signals (De Falco et al., 2024), but also in regulating consciousness, attention, sleep and motor control (Redinbaugh et al., 2020; Zhou et al., 2021). These tasks typically involve one or more thalamic neuronal clusters, named subthalamic nuclei. For instance, the lateral geniculate nucleus is involved in visual processing and the medial geniculate nuclei in auditory processing. (Ghodrati et al., 2017; Winer, 1984)

The thalamus’ broad involvement in neural circuits makes it a critical region for understanding both normal brain function and a variety of neurological and psychiatric disorders. Dysfunctions in the thalamus have been implicated in several conditions, including psychosis, epilepsy, and Parkinson’s disease. For example, structural and functional abnormalities in the thalamus have been reported in schizophrenia, particularly affecting cognition and sensory processing (Andreasen et al., 1994; Byne et al., 2009; Delavari et al., 2024). Additionally, thalamic degeneration is a hallmark in neurodegenerative disorders like Parkinson’s disease, where the loss of dopaminergic neurons impacts motor control (Braak et al., 2004).

The neonatal period represents a critical window in human brain development, where rapid and complex neurodevelopmental processes lay the foundation for future cognitive and motor functions (Bhat et al., 2014; L. Cai et al., 2024; Godfrey & Barker, 2001; O’Donnell & Meaney, 2017; Volpe, 2009). During this phase, the brain undergoes extensive synaptogenesis, myelination, and structural reorganization. The thalamus, in particular, is a vital structure during this stage, as it plays an essential role in the early formation of sensory and cognitive networks. In fact, one of the first fibre bundles to form in the human brain are the thalamocortical tracts that reach completely all the neocortical areas by around week 30 of gestation. Delineating thalamic nuclei has been typically performed using histology (Morel et al., 1997). However, as of today and to the best of our knowledge, no neonatal histological atlas exists. Nevertheless, recent advancements in magnetic resonance imaging (MRI) have allowed in vivo thalamic parcellation (Segobin et al., 2024).

Current structural MR imaging provides detailed anatomical information, including neonatal thalamic volume and shape development (Li et al., 2024). However, conventional T2-w and T1-w contrasts provide insufficient information (Ferradal et al., 2019; Toulmin et al., 2015) for thalamic nuclei segmentation. While advanced MRI contrast are being developed for adults (Martinez et al., 2024; Segobin et al., 2024), their extension to infants is not straightforward, given the limit in acquisition time and subject motion. Alternatively, diffusion MRI captures the brain’s microstructural properties, including white matter tracts connectivity patterns (Eaton-Rosen et al., 2014; Jaimes et al., 2018; Jakab et al., 2020; H. Ji, Payette, et al., 2024; H. Ji, Wu, et al., 2024; Oldham et al., 2024; Zheng et al., 2023) in normal or abnormal (e.g. pre-term birth and congenital heart disease) conditions.

Functional MRI (fMRI), on the other hand, offers insights into brain activity by measuring blood oxygenation level-dependent (BOLD) signals, which are indirect markers of neuronal activity. fMRI, in particular, has been crucial in characterizing developing brain thalamocortical functional connectivity (Alcauter et al., 2014; Y. Cai et al., 2017; Ferradal et al., 2019; Fransson et al., 2009; Toulmin et al., 2015, 2021).

This connectivity pattern has been used to parcellate the neonatal thalamus into different nuclei. For instance, Alcauter et al., 2014 aimed to characterize development, between neonates, 1- and 2- year old infants, including cognitive correlates. The neonatal cohort included 112 subjects. The authors used a winner-takes-all (WTA) approach on the highest group-level partial correlation of BOLD signal with 9 cortical areas (sensorimotor; auditory; medial visual; occipital poles; lateral visual; default mode; salience; right and left frontoparietal). They found that thalamus–sensorimotor and thalamus–salience networks are the most present at term, while thalamus–medial visual and thalamus–default mode networks are not evident until 1 year of age. They also showed that the sensorimotor cluster covers both visual and auditory areas, suggesting reduced specialization at the neonatal age.

Another study (Y. Cai et al., 2017), in which the target networks were derived based on the neonatal Automated Anatomical Labeling (AAL) template (Shi et al., 2011), aimed at comparing thalamocortical connectivity between term (N=31), pre-terms (N=22) and pre-terms with white matter lesions (N=22). The authors reached similar conclusions as Alcauter et al., 2014 where no significant thalamic-DMN (Default Mode Network) connection was observed, as opposed to sensory-motor and salience networks that were the most connected to thalamus. The medial part of the thalamus was connected to the latter while the dominant lateral part was connected to the former. The authors also showed that pre-terms with white matter lesions exhibit an altered thalamo-SA (Salience) connectivity compared to term subjects. This study utilized a WTA segmentation approach to the 7 cortical networks, namely sensorimotor, auditory, medial visual, default mode, salience, left and right frontoparietal networks.

A different study (Toulmin et al., 2015) included very pre-terms (N=43) and term (N=23) subjects, both scanned at term age, to parcellate the thalamus. Similar to Alcauter et al. (2014) and Y. Cai et al. (2017), the authors used a WTA segmentation approach on the highest correlations to 9 cortical regions (primary sensory motor, primary auditory, sensory motor association, primary visual, temporal, prefrontal, lateral parietal, frontoparietal insular, and anterior cingulate). However, preprocessed BOLD signal from all subjects was concatenated (and not z-scored as in Alcauter et al., 2014; Y. Cai et al., 2017) in the temporal dimension and resting state components (i.e. seed regions) were defined using ICA (Independent Component Analysis) with a 25 dimensionality. Each thalamic voxel was then assigned to a resting state component depending on which network had the highest z-score at that voxel, and 9 cortical regions were extracted. Overlapping thalamic connectivity profiles that are correlated with multiple cortical regions were identified. This suggests as in Alcauter et al., 2014, a lack of specialization of thalamic nuclei at a neonatal age. The study also found that extreme prematurity is associated with decreased connectivity between the thalamus and prefrontal, insular and anterior cingulate cortex, and increased connectivity with the lateral primary sensory cortex. The same authors followed (Toulmin et al., 2021) a pre-term cohort of 102 subjects at 2 years age to predict motor and cognitive outcomes based on thalamocortical connectivity at neonatal term age. A similar approach was employed as in Toulmin et al., 2015, where this time 11 cortical areas with bilateral hemispheric representations were used for analysis. Their results show a largely symmetrical representation of these cortical areas on the thalamus, at the exception of the lateral prefrontal cortex. Highly overlapping clusters were also observed with dominance of the motor cluster spanning all the thalamus. They found that pre-motor association cortex connectivity to thalamus correlates with motor function, while primary sensorimotor cortex and thalamus connections correlate with cognitive scores.

Finally, Ferradal et al., 2019 aimed at analyzing the interactions between functional (fMRI, 350 timepoints) and structural (diffusion MRI, 60 gradient directions) thalamocortical connectivity in 20 healthy full-term neonates (mean age at scan 42.7 weeks). For fMRI clustering, using 7 pre-defined cortical regions (prefrontal, premotor, primary motor, somatosensory, posterior parietal, occipital and temporal), partial correlations were computed between the mean cortical signals and the time-series of each thalamic voxel. After z-scoring and averaging, each voxel was assigned to the highest cortical region it was connected to (i.e. WTA). The same cortical areas were used as target regions of fiber tracts extracted from the diffusion MRI signal. Their results show a significant overlap between functional and structural thalamic parcellations, especially in primary sensory areas, as opposite to higher-order association areas such as temporal and posterior parietal cortices. The functional parcellation is highly dominated by the somatosensory cluster, while the pre-frontal cluster is situated near the mediodorsal area of the thalamus, whereas the temporal cortex shows a cluster located around the pulvinar nucleus.

While these studies are highly important in showing the trends of identifying functional subdivisions of the thalamus based on the connectivity patterns to the cortex, both for full- and pre-term neonates, they remain limited in the number of subjects (20-112) and in the number of timepoints (150-350) of the fMRI acquisitions. Moreover, the neonatal brain’s process of development and specialization (Alcauter et al., 2014; Toulmin et al., 2015, 2021) is in its early stage. Additionally, the thalamus is involved in a variety of tasks, including integrative functions (Greene et al., 2020). These two points suggest that the appropriate approach to parcellate the neonatal thalamus should take into account the complexity and variability of its connections to other brain regions. Thus, a thresholding approach such as the winner-takes-all as employed by all previously described studies might not be suitable to capture this particularity of thalamic nuclei connectivity patterns. Finally, the choice of the number of thalamic clusters is highly heterogeneous and is rigidly linked (except Toulmin et al., 2015, 2021) to the few pre-defined cortical seeds. In fact, methods employing fine grained clustering algorithms on functional connectivity based quantities to parcellate a specific brain regions (Delavari et al., 2024; Garcea & Mahon, 2014; B. Ji et al., 2016; Luo et al., 2020; Ren et al., 2019) have never been applied in the case of developing brains, to the best of our knowledge.

In this study, we leverage the large neonatal research cohort of the developing Human Connectome Project (dHCP), using more than 700 subjects, acquired at 2300 temporal volumes, to parcellate the thalamus between 37 and 44 weeks of post-menstrual age using an introduced framework: KNIT (K-means for Nuclei in Infant Thalamus). We employ a segmentation approach based on the connectivity pattern of each thalamic voxel to 40 brain regions, both cortical and subcortical areas, where we aimed to characterize brain thalamic nuclei development. We also determine the number of clusters using bootstrapping strategies, from which we derive uncertainty maps to assess the voxel-wise reliability of the thalamic clusters. Finally, we compare the advantages and limitations of this approach with the conventional winner-takes-all method.

## 2 Methods

### 2.1 Data

#### 2.1.1 Developing Human Connectome Project (dHCP) protocol

The third data release of the publicly available dHCP dataset (Edwards et al., 2022)^1^ was used in this study. We selected 730 neonatal dHCP subjects with postmenstrual age at scan [36.57, 44.43] weeks (mean±std at birth = 38.16±3.9 weeks). Subjects were divided into age groups according to Table 1. Data were obtained using a 3T Philips Achieva MRI scanner equipped with a specialized neonatal imaging setup, which included a 32-channel head coil. Resting state fMRI (rs-fMRI) data were acquired using a multislice gradient-echo EPI sequence with multiband excitation (TE=38 ms; TR=392 ms, Multiband factor=9, Flip Angle=34°). 2300 timepoints of Blood Oxygenation Level Dependent (BOLD) signal image were acquired at an isotropic resolution of 2.15 mm. Single-band EPI reference scans were also acquired with bandwidth-matched readout, along with additional spin-echo EPI acquisitions. In order to correct for EPI-induced distortions, field maps were obtained from an interleaved spoiled gradient-echo sequence. Anatomical T1- and T2-weighted images were also acquired during the same scan session, with a spatial resolution of isotropic 0.8 mm using Fast Spin Echo and Inversion Recovery Fast Spin Echo, respectively. Using all the above-mentioned images, pre-processed 4D rs-fMRI volumes (Fitzgibbon et al., 2020) (for motion and distortion correction (Andersson et al., 2018), high-pass filtering and denoising) were provided and were hereby used in our analyses. T1- and T2-weighted age templates were also provided (Makropoulos et al., 2018). Further information about data acquisition and preprocessing can be found in Fitzgibbon et al., 2020.

**Table 1:** Age groups and their number of subjects from the dHCP dataset (Edwards et al., 2022).

| Age (weeks) | Number of Subjects |
| --- | --- |
| 37 | 32 |
| 38 | 46 |
| 39 | 70 |
| 40 | 104 |
| 41 | 153 |
| 42 | 135 |
| 43 | 103 |
| 44 | 87 |

#### 2.1.2 Brain parcellation labels

With the T1-w and T2-w images, label masks of 87 brain region were provided (Makropoulos et al., 2018). Thalamic labels were unified from the two hemispheres to define the thalamus mask. Moreover, 40 regions of interests (ROIs) were extracted from these labels (details in Table 2 of Supplementary Materials), namely cortical areas, deep brain areas (hippocampus, amygdala, parahippocampal gyrus and caudate nucleus) and the cerebellum (Figure 1 top left for visualization).

**Figure 1:**
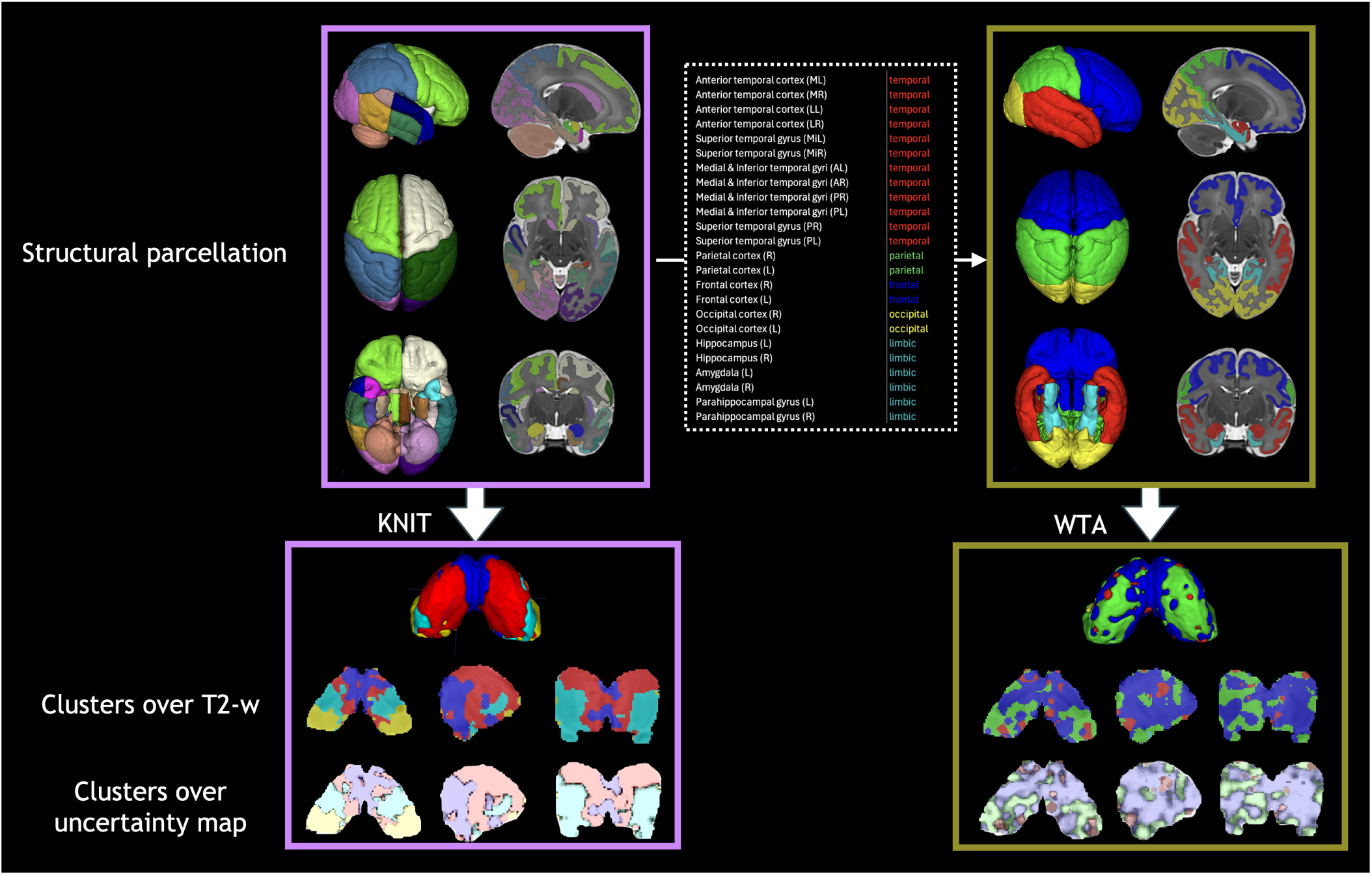
KNIT (k=5 in k-means) using 40 ROIs overlayed on T2 maps (detailed in Supplementary Table 2) a of 41 weeks neonates (N=153) compared to the WTA approach, where the ROIs merging is shown on the top middle box. Clusters are overlayed on T2-w age-matched template. The bottom row shows the same clusters overlayed on the uncertainty maps (as defined in Section 2.3.1), showing a high uncertainty at the cluster borders (gray scale reversed for better visualization). All images including isosurfaces (i.e. 3D rendering) were created using ITKSNAP (Yushkevich et al., 2016).

### 2.2 Thalamic functional connectivity (FC) pipeline

First, the label maps in the T2-w space were registered using SITK (Beare et al., 2018) to the BOLD images using an affine transformations. In the subject native space, functional connectivity is computed between each thalamic voxel and the 40 ROIs defined above. The first 5 timepoints of the BOLD signal were discarded to allow the signal to approach steady state (as in Hu et al., 2022. We performed spatial smoothing with a 3 mm full-width-at-half-maximum (FWHM) Gaussian kernel and temporal low-pass filtering (0.08 Hz low-pass cutoff) on the pre-processed dHCP rs-fMRI before computing FC maps. Each subject FC in the native fMRI space is registered and resampled to the age-matched T2-w template space of the thalamus. For instance, a subject of 39.8 weeks will be registered to the 40 weeks template. Specifically, a rigid registration followed by an affine registration with mutual information as the cost function are applied. Before averaging FC maps, all resampled subjects to the template space were merged together and converted to approach a normal distribution using the Fisher-Z transform (Fisher, 1915).

### 2.3 KNIT versus Winner-Takes-All (WTA) parcellation

In the proposed method, we use all the parcellation information by computing the functional connectivity using the correlation between each thalamic voxel and 40 brain regions (parcellation described in Supplementary Table 2 with respective abbreviations). Each vector, associated to each thalamic voxel, is clustered based on its 40 components that represents its connectivity profile to the 40 brain regions. Clustering is performed using k-means with a *cosine similarity* distance with *k* ∈ {2, 3, 4, 5, 6, 7} clusters. The Hungarian algorithm (Kuhn, 1955) is then applied to match the cluster IDs between the different subjects bootstraps (described in Section 2.3.1 below) and age groups (with the first bootstrap of the 40 weeks age group as a reference).

In the WTA method (Alcauter et al. (2014), Y. Cai et al. (2017), Ferradal et al. (2019), Fransson et al. (2009), and Toulmin et al. (2015, 2021)), for each thalamic voxel, we compute the functional connectivity (FC) using the partial correlation between that voxel and each of the four lobes (plus limbic areas) (details and visualization of the merged labels in Figure 1, top), and the maximum value among the 5 values is assigned to one lobe/cluster. Using this winner-takes-all strategy, each voxel is exclusively assigned to temporal, parietal, frontal, occipital or limbic clusters. The final clusters are the result of the majority voting among the different bootstraps (described in Section 2.3.1 below).

As a final step for both methods, we post-processed images to assign voxels of small (less than 16 contiguous voxels, i.e. connected components) clusters found inside or highly surrounded by a dominant cluster, to that dominant cluster.

#### 2.3.1 Bootstrapping and uncertainty quantification

Two types of bootstrapping were applied: (i) *subjects bootstrap*: a random selection of 75% from all the subjects among each post-menstrual age group (40 bootstraps); (ii) *timepoints bootstrap*: a random selection of 75% BOLD timepoints for a fixed number of 25 subjects for each post-menstrual age group (10 bootstraps). These computations help in choosing an optimal number of clusters *k*, on pinpointing more uncertain voxels and clusters and on generating the final clusters that were taken based on a majority vote among the 40 subjects bootstraps. An *uncertainty* map was also computed, based on the subjects bootstraps, from the inverse of the proportion of the majority label on each voxel *U* (*v*). The average uncertainty across voxels and clusters, normalized by the total subject number of the corresponding age group (*U*), is also reported:

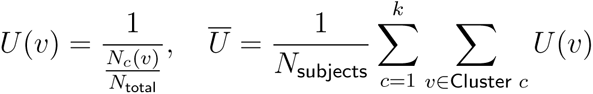

*N_c_*(*v*) is the number of times voxel *v* is assigned to cluster *c* and *N*_total_ is the number of bootstraps (40).

In order to decide on an optimal number of clusters, variability across the different bootstraps was computed using the Coefficient of Variation on the Intersection over Union (CoV of IoU) over every segmentation pair of bootstraps. This was performed both in subjects (i) and the timepoints (ii) bootstraps. The timepoints bootstraps for a fixed number of subjects also help us compare the variability across age groups, by removing the confound of the number of subjects (i.e. age groups containing more subjects will have an intrinsic lower variability). We have also computed the Silhouette score, i.e. the proximity between data points and there assigned cluster compared to other clusters (the higher the better) for *k* ∈ {2, 3, 4, 5, 6, 7} clusters.

#### 2.3.2 Functional connectivity analysis

After computing the different clusters, we looked back at the FC maps that correspond to each cluster in order to analyze the connectivity pattern of that cluster. We also computed the z-score of the FC maps of each cluster and considered regions that fall above +1 or below -1 as significantly connected or disconnected, respectively, to that cluster. To facilitate analysis and if right and left connectivities are similar, FC maps were bilaterally merged in the z-score plots.

Specifically, for each ROI, we take the median FC value in each cluster and in each bootstrap and average across bootstraps to get the final conectivity pattern for this specific cluster.

### 2.4 Across age metrics

For each age group we have computed several metrics to analyze developmental trends:

#### Bootstraps and uncertainty

Timepoints bootstraps and normalized uncertainty based on the subjects bootstraps as described in Subsection 2.3.1.

#### Relative cluster volumes

The relative volume of each cluster is defined as the volume of the cluster divided by the total thalamic volume. Mean and standard deviation are also reported across the different subjects bootstraps.

#### Clustering symmetry index

It is aimed to quantify the bilateral symmetry of the majority voted clusters. It is computed by mirroring the left hemisphere clusters to the right hemisphere ones and computing the agreement between the superposed voxels, i.e. 1 if similar clusters and 0 otherwise. The right hemisphere is also mirrored to the left one and the agreement is computed. An average of indices, divided by the average of the total number of voxels in each hemisphere, is then taken as the final index (0 as completely asymetrical and 1 as completely symmetrical).

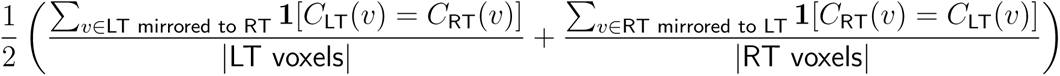

refers to the cluster ID of voxel *v* in the right/left thalamus (*X* ∈ {*R, L*}).

#### Ipsilateral and contralateral connectivity symmetry

Here we quantify how much is the right hemisphere thalamus connected to the right hemisphere ROIs and to the left hemisphere ROIs, and vice versa, using the majority voted clusters. For the ipsilateral connectivity index, it is defined as the ratio between the average connectivity of the right thalamus to the right hemisphere and the average connectivity of the left thalamus to the left hemisphere. Inversely, the contralateral connectivity index is defined as the ratio between the average connectivity of the right thalamus to the left hemisphere and the average connectivity of the left thalamus to the right hemisphere.

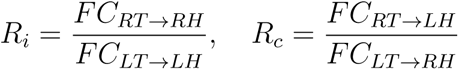

*FC_XT_*_→*Y*_ *_H_* refers to the functional connectivity of the right/left thalamus to the right/left hemisphere (*X, Y* ∈ {*R, L*}).

#### Global thalamic connectivity

It is computed by averaging the absolute value of functional connectivity of each thalamic voxel to each of the 40 ROIs and then bilaterally averaging the values between the two hemispheres. No bootstrapping was applied here and all the subjects from all age groups were included in the computation.

#### ROI-ROI connectivity

Functional connectivity matrices between each pair of the 40 ROIs. They are calculated using the Pearson correlation of the normalized averaged time series, using 25 subjects (average FC) in each age group. In addition, we calculate the differences between the FC matrices of consecutive age groups (i.e. 44 and 43 weeks, 43 and 42 weeks, etc.). Finally, we compute a global ROI-ROI connectivity by taking the average absolute values in each matrix to assess the global brain connectivity variations across development.

## 3 Results

Qualitative results thalamic parcellations show a physical proximity and spatial coherence of voxels belonging to the same clusters as well as a symmetrical topographical representation across hemispheres as depicted in the bottom left of Figure 1. It is worth noting that these two properties were obtained with a voxel-wise clustering without any local spatial prior (e.g. spatial coordinates information), i.e. in a completely unsupervised manner. We start by analyzing the thalamic nuclei from the oldest group as this is the more mature state across our age span, followed by a developmental analysis.

### 3.1 Number of clusters selection

We aim to set the number of clusters *k* of k-means equal for all age groups in order to analyze their evolution across development. We thus conduct a bootstrapping analysis to determine the optimal *k*. Figure 2a shows for the 44 weeks group, the variability (mean CoV of IoU) of the clusters overlap for each cluster number *k*, for the timepoints (a) and the subjects (b) bootstrapping strategy (other age groups in Supplementary Figures 11, 12). We observe, as expected, a general trend of increased variability with an increased number of clusters. However, we often observe a higher gap, especially when the number of subjects is high (i.e. age groups of 40, 41, 42 and 43 weeks), from 4 to 5 and 5 to 6 clusters. If we compare the variability of the timepoints bootstrapping across age (Figure 2b, left) for a fixed number of subjects (to remove the number of subjects confound), we observe a clear evidence that variability reduces with age, meaning thalamic nuclei parcellation can be more robustly performed for older neonates with a high number of clusters, suggesting more specialization and differentiation across age. The Silhouette score shows different trends (Supplementary Figure 13) for different age groups with a general low standard deviation for the different (subject) bootstraps for the cluster number *k* = 5. Given the three experiments above and their general hints, we have opted for *k* = 5 (more details on the selection procedure in Supplementary Figure 14).

**Figure 2:**
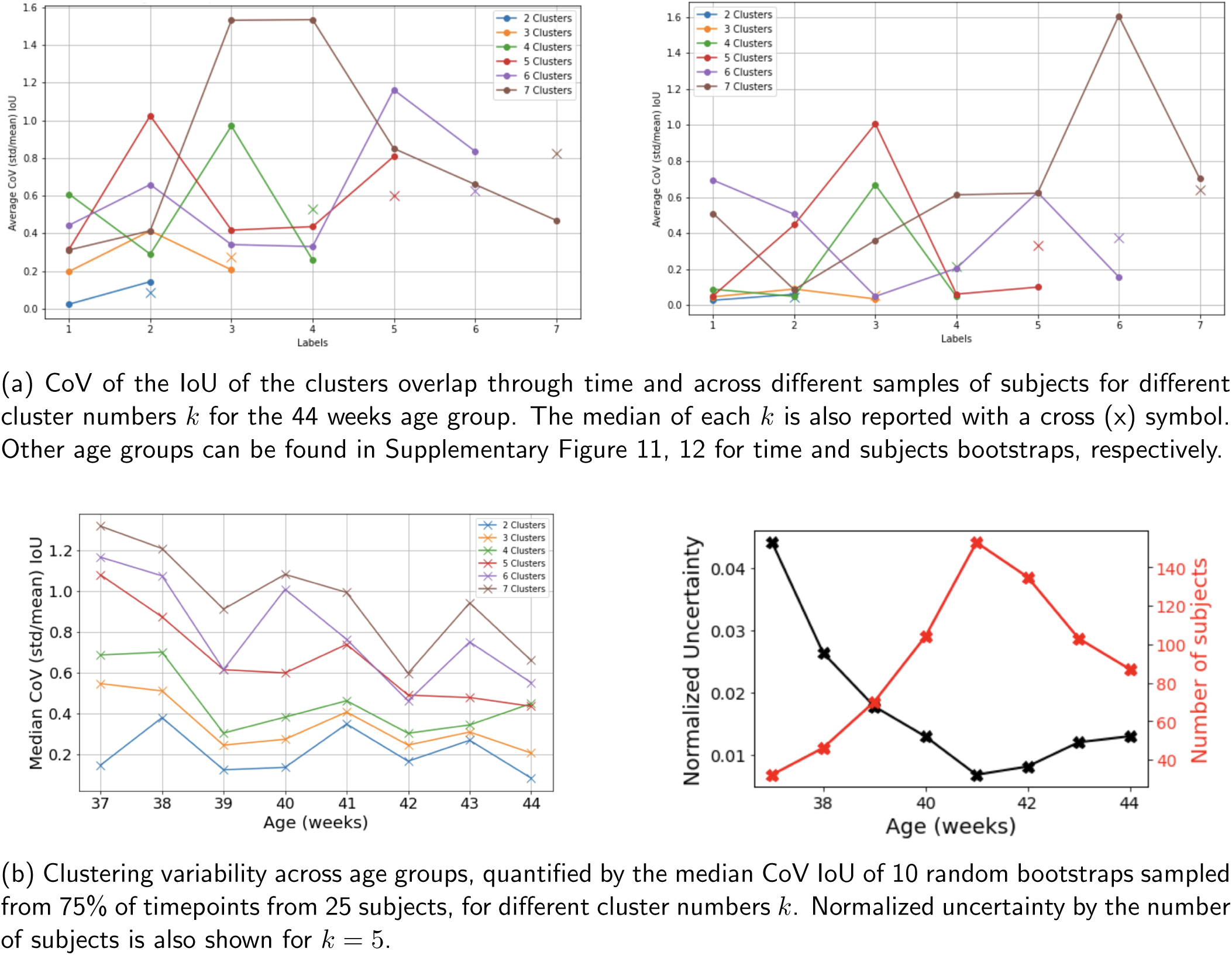
Analysis of clustering variability: (a) top panel shows the 44 weeks variability analysis; (b) bottom panel presents the variability analysis across age groups.

### 3.2 Cluster analysis at 44 weeks

We present first the functional thalamic cluster analysis in the oldest age group as this is the more mature state across our age span. The connection to findings of the literature on neonatal thalamic parcellations will be detailed in the Discussion section.

Cluster 1 represents the bilateral posterior thalamus, grossly overlapping with the pulvinar and the geniculate nuclei. When looking at the connections of this cluster (Figure 3), we find the strongest connection to be to the temporal gyrus (covering superior, middle and inferior temporal gyrus with a predominance on the right side), occipital lobe and parietal lobe. This finding is interesting because the pulvinar area of the thalamus is known to take part in cognitive processing of the audio visual sensory stimuli. Additionally, the geniculate nuclei send direct connections to the auditory cortex (superior temporal gyrus) and the visual cortex (central occipital lobe). Moreover, the parietal lobe is known to be a major hub for sensory processing, integration and comprehension.

**Figure 3:**
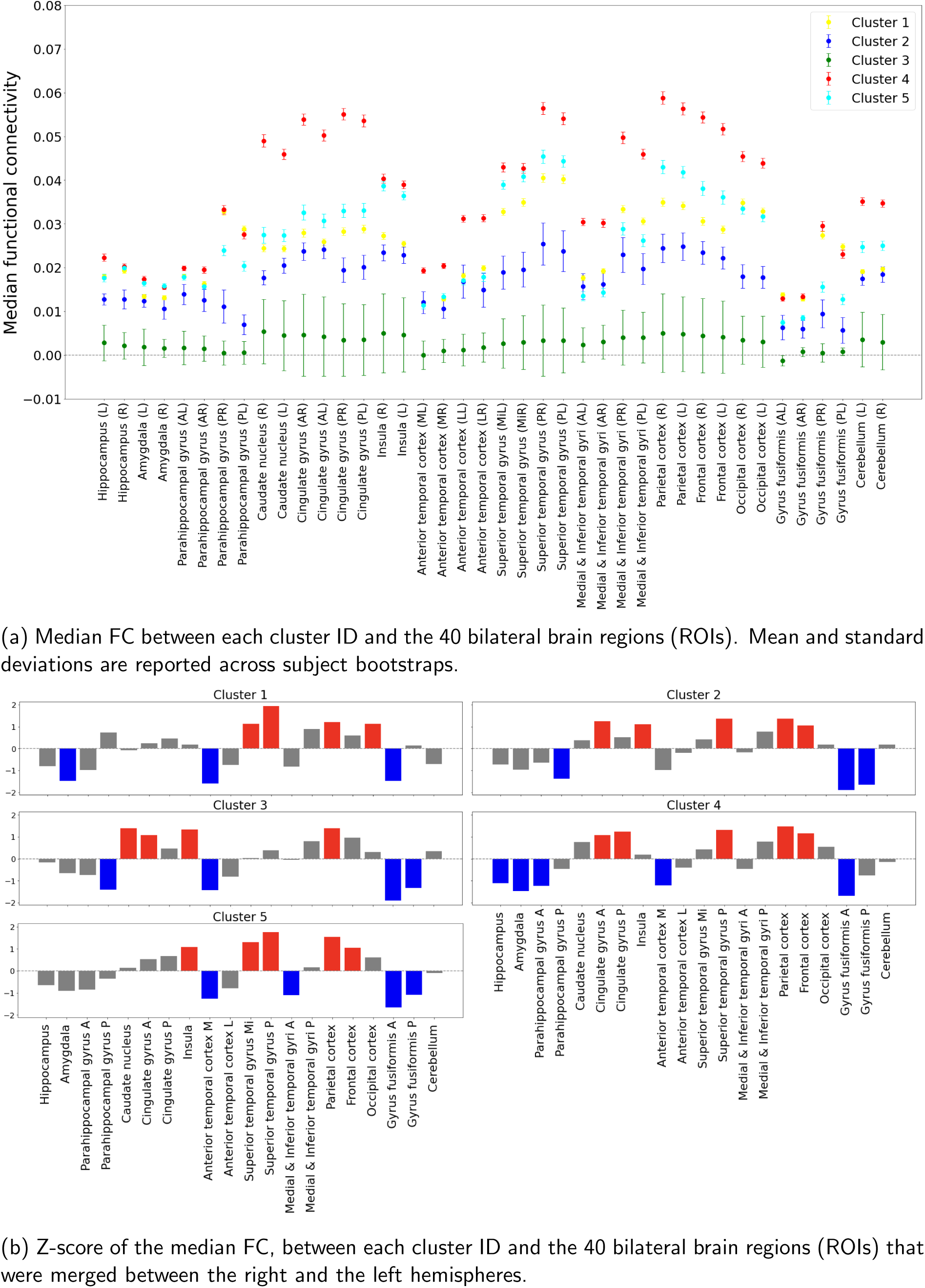
Median functional connectivity (FC) between each cluster ID and the different ROIS, for the 44 weeks age group. Other age groups can be found in Supplementary Figure 16.

Cluster 2 represents the medial part of the thalamus. Overall connectivity profile of this cluster is similar to cluster 1 in high values of connectivity with the posterior part of superior temporal gyrus and the parietal lobe. However this cluster additionally connects to the frontal, bilateral insula and the anterior cingulate cortex. The last 3 are famously known to form the salience network, for which the function is defined as directed attention and higher order cognition. Interestingly, the medial part of the thalamus is known for its role in executive functions (such as planning, cognitive control, working memory, etc.) which also famously involve the salience network.

Cluster 3 demonstrates low connectivity to majority of the regions with confidence interval crossing 0. Cluster 3 also contains the least amount of voxels with an asymmetric localization within the thalamus. Overall, we believe this cluster represents a small sample of voxels with high noise and therefore is a discarded cluster. It is worth mentioning that this cluster also appeared when the number of clusters *k* in k-means was set to 4 or 6 for example.

Cluster 4 is the largest cluster covering the dorsal, and anterior part of the thalamus, with a connectivity profile covering the frontal cortex, superior temporal gyrus, and parietal lobe. This cluster spatially overlaps with the dorsal thalamus, which is known for strong reciprocal connections with the prefrontal cortex, a key component of the frontoparietal network (FPN).

The dorsal nucleus plays a central role in cognitive processes like decision-making and executive functioning, supporting the FPN in maintaining and manipulating information for goal-directed behavior. The overlap of the connectivity profile of the Cluster 4 with the FPN suggests its involvement in integrating inputs across cortical regions to facilitate adaptive cognitive control.

Cluster 5 is located in ventrolateral areas of the thalamus that cover the nuclei of the thalamus known as the motor thalamus. In terms of connectivity, the cluster 5 connects to parietal and superior temporal gyrus, additionally it connects to frontal and insular cortex. In that sense, it has a similar connectivity profile to cluster 2, except that cluster 2 has a lower connectivity to limbic areas compared cluster 5, making the distinction between the two clusters possible within the clustering algorithm.

Unfortunately, within our cortical segmentation, motor regions were included as part of the frontal lobe, therefore, we did not expect the motor regions of the thalamus to be represented within our segmentation. However, potentially, the combination of connections in the connectivity profiles could give rise to a additional information, resulting in a more fine grained parcellation. This point is further discussed in the discussion.

### 3.3 Comparison of KNIT and WTA: across age analysis

Figure 4 and Figure 5 show qualitatively the development of the thalamic nuclei from 37 to 44 weeks of age for KNIT and the WTA approach, respectively. We can also visually assess the strength of the first method that is further explored below, along with WTA for certain experiments.

**Figure 4:**
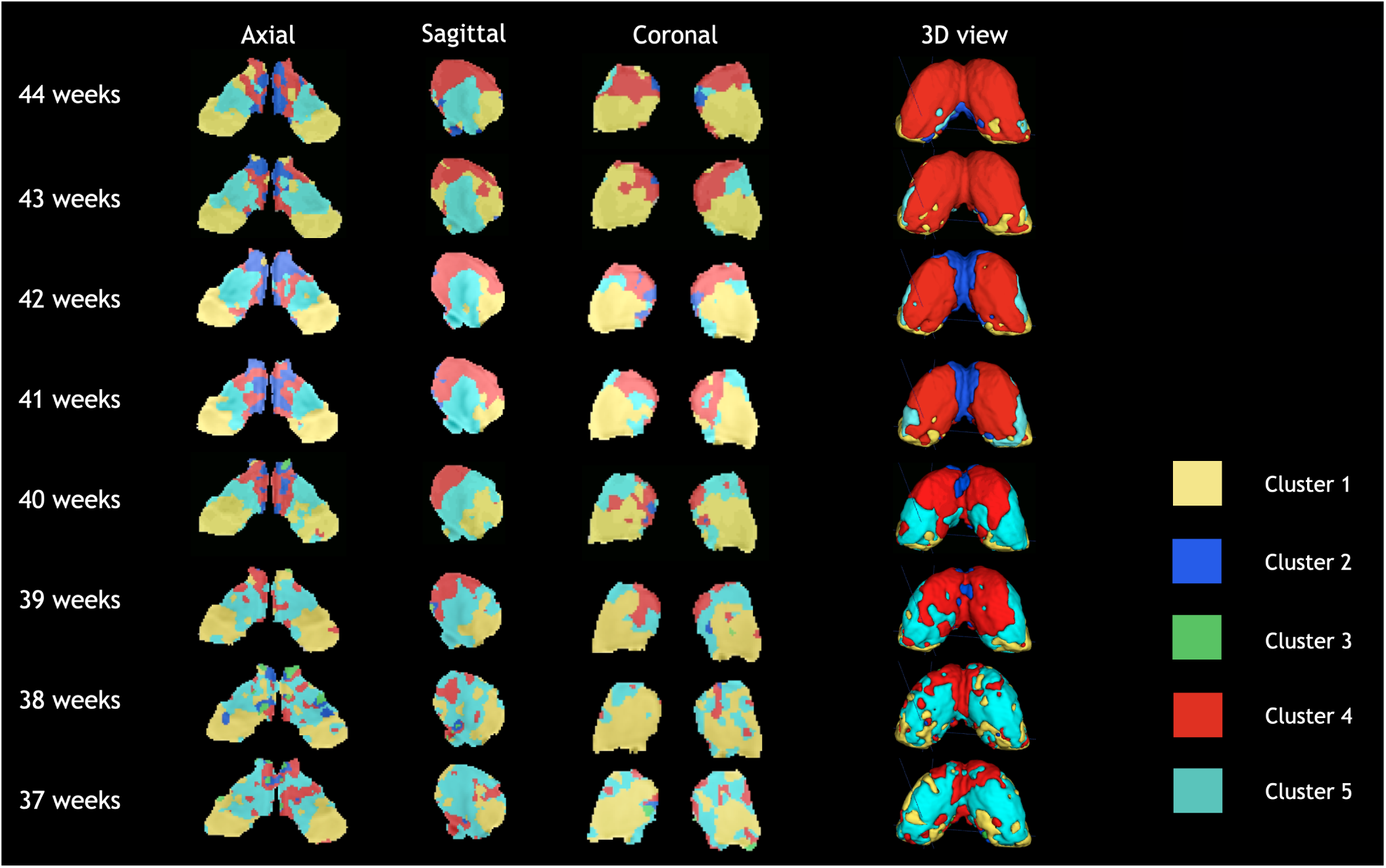
Orthogonal views and 3D isosurfaces of the group-level clusters evolution across the age span for KNIT (*k* = 5 in k-means). Right hemisphere corresponds to left part in the axial and coronal views.

**Figure 5:**
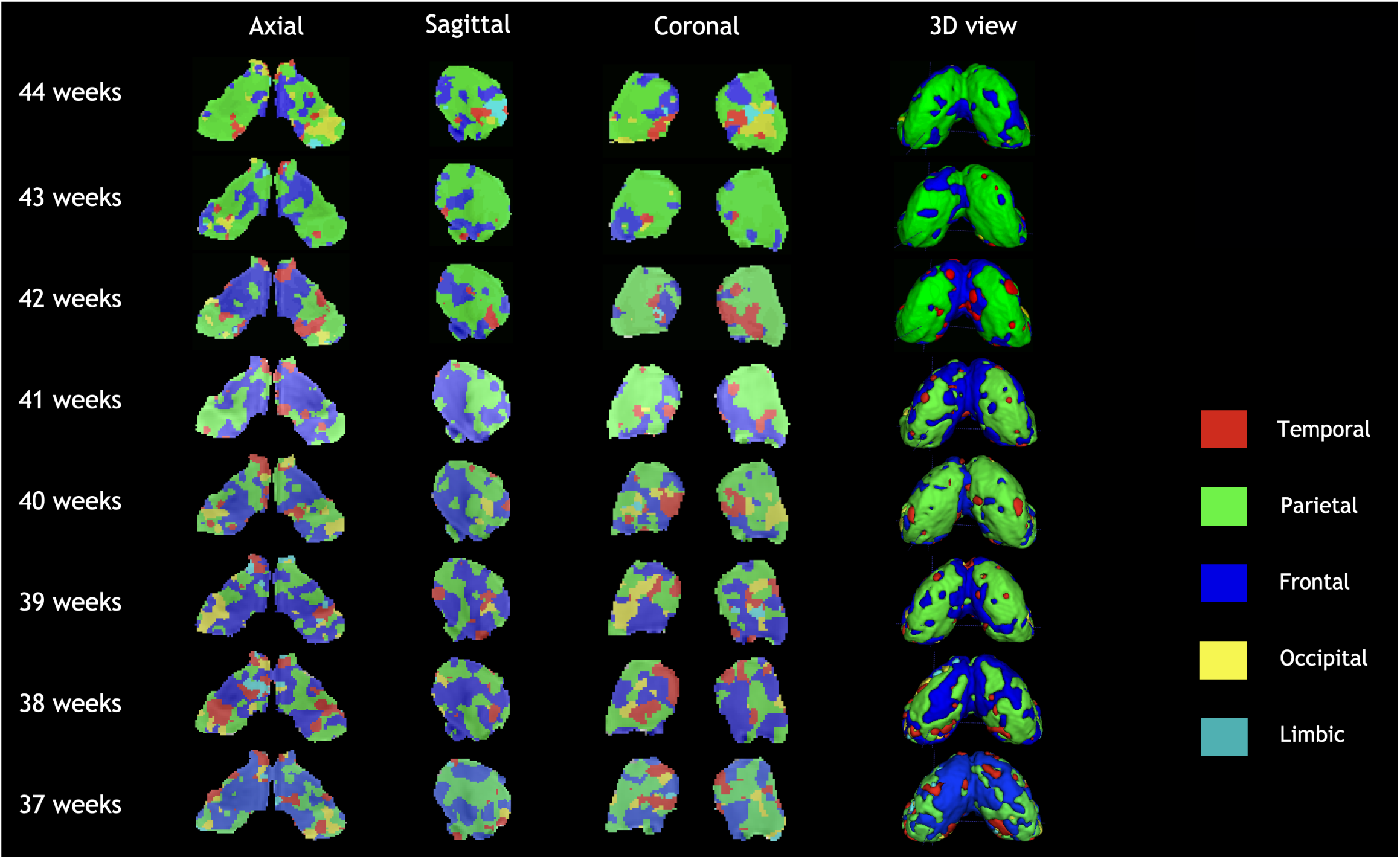
Orthogonal views and 3D isosurfaces of the group-level clusters evolution across the age span for WTA approach. The right hemisphere corresponds to left part in the axial and coronal views.

#### Relative cluster volumes and thalamic connectivity

Here we aimed to assess the trajectory of the volume (Figure 6) and the intensity of the average connection of each thalamic segments across age. We observed that the most strong change of relative volume occurs in form of decreasing volume of the ventrolateral cluster (cluster 5) and increasing volume of dorsomedial cluster (cluster 4). Additionally the intensity of the connectivity of the cluster 4 increases over age, overpassing cluster 5 as 41 weeks. This is in line with translational evidence showing the development of major afferent connections of the mediodorsal thalamus to the cortex happening postnatally and stabilizing in the infantile years (Rios & Villalobos, 2004). Additionally, the pulvinar cluster (cluster 1) containing audiovisual nuclei of thalamus also shows a weak decrease of relative volume. As the dorsomedial thalamus is implicated in networks invloving higher order functions whereas ventrolateral is more involved in motor processing. This shift of relative volume and connectivity strength could be indicative of a shift towards formation of higher order nuclei of thalamus. It is worth mentioning that cluster 2 volumes at age 38 and 37 weeks is extremely low and this is accompanied by a disruption in the connectivity pattern compared to other age groups (Supplementary Figure 16). This might also be explained by the smaller sample size of these two age groups.

**Figure 6:**
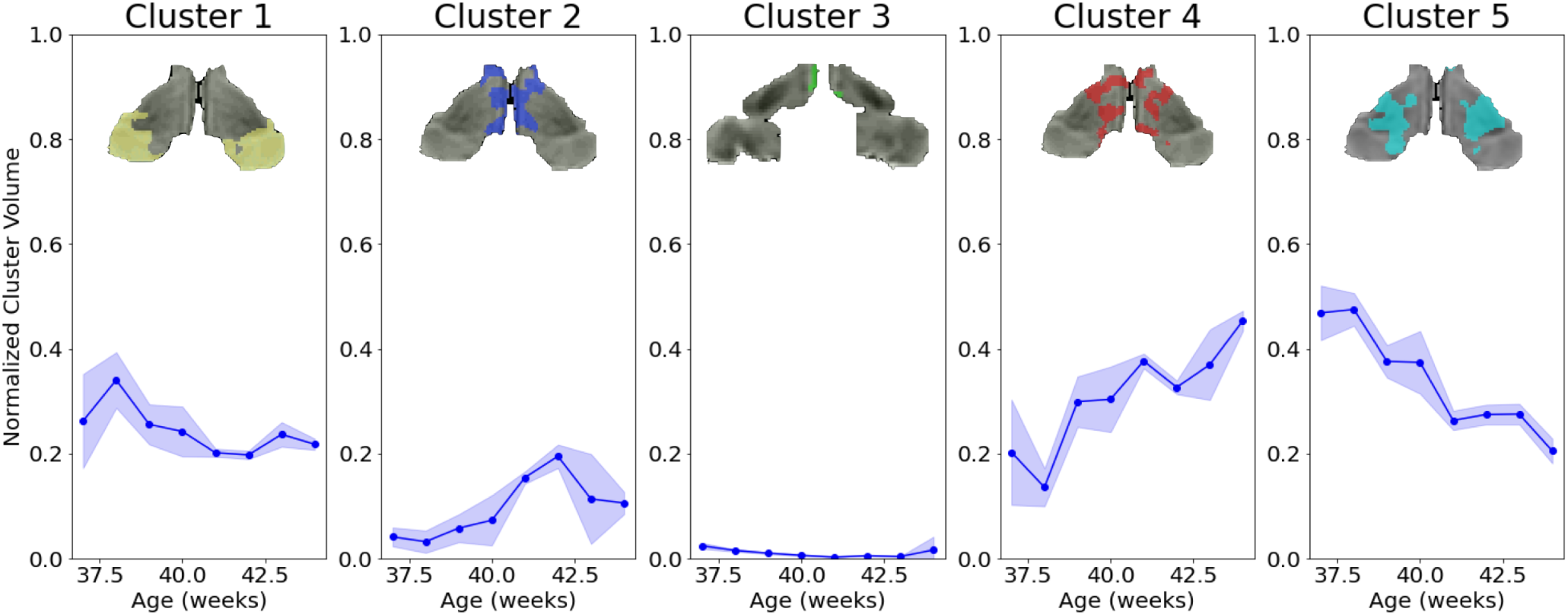
Developmental evolution of the normalized volumes (across whole thalamic volume) for each cluster. Mean and standard deviations across subject bootstraps are displayed.

For the WTA approach, we mainly observe two dominant clusters, representing more than 85% of the thalamic voxels (Figure 15). Namely, a cluster that is highly connected to the frontal cortex, that would grossly correspond to a combination of cluster 2 and cluster 5 in KNIT; and a cluster that is highly connected to the parietal cortex that would grossly correspond to cluster 4. It is worth to mention that that these correspondences are approximative as no clear mapping can be extracted given the high connectivity of all voxels to several brain areas (i.e the lack of specialization).

In terms of volume change across age, the parietal cluster is growing in age whereas the frontal one is shrinking. This can be explained by a growing involvement of somatosensory areas relative to motor areas. The occipital cluster, that represent between 1% and 8% of the clusters volume, is mostly situated in the ventrolateral area. The temporal cluster does not show a specific spatial preference, and the limbic cluster is extremely small .

Clearly, the WTA approach does not allow enough granularity to delineate clusters based on their complex connectivity pattern, as particularly at this age, voxels are connected to many dominant areas and not specifically to one, as per se forced by this method. As an example, the pulvinar cluster that was delineated by KNIT, was completely missed by the WTA method.

#### Uncertainty estimation

Figure 2b (right) shows a decrease in the normalized uncertainty by the number of subjects through age. In fact the slight increase after 41 weeks is likely due to the rapid decrease in the number of subjects, as shown between the absolute slopes of the two curves. In other words, the decrease in uncertainty will very much likely have continued if the number of subjects was higher between 42 and 44 weeks groups. This drop is also confirmed by the decreasing variability through age for the timepoints bootstrapping method using a fixed number of subjects (Figure 2b, left). Therefore, these results could hint towards an increase of functional specialization of the thalamus across the age span.

#### Clustering symmetry index

We observe (Figure 7a) an overall general bilateral symmetry of the k-means clustering, that also seem to increase with age, which can partly be due to the increased number of subjects for certain age groups. However, the drastic decrease in number of subjects from 42 to 44 weeks, cannot solely explain the non-decrease of the symmetry index. This suggests a general increased symmetry through age. Moreover we also observe that the dorsal thalamus (cluster 4) becomes more symmetrical with age, which can be explained by its gain in volume.

**Figure 7:**
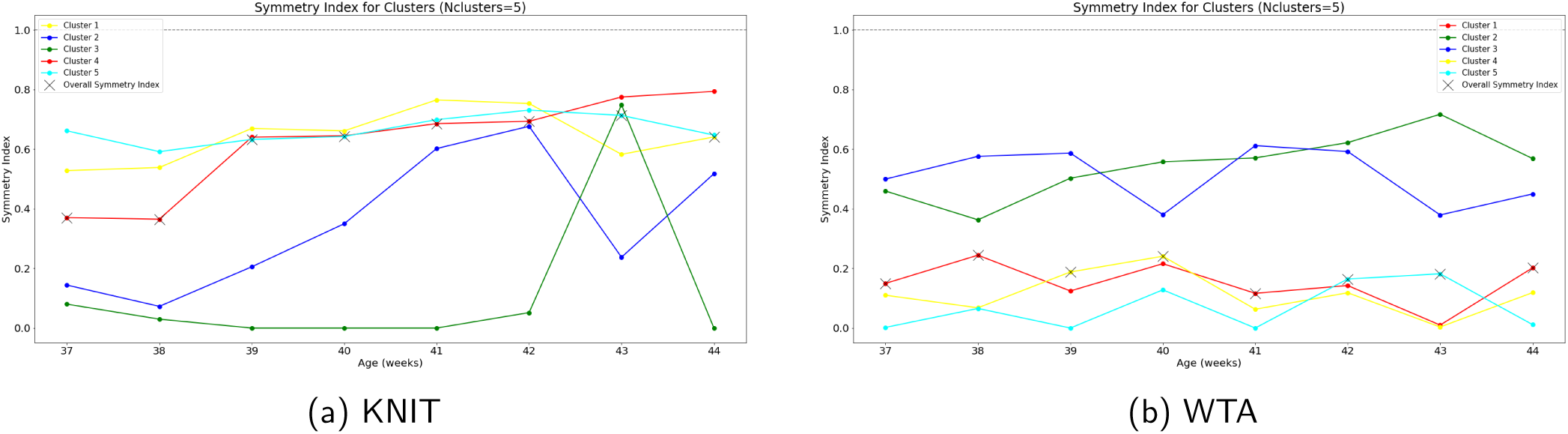
Bilateral clustering symmetry index of the thalamic labels as defined in Section 2.4. The median symmetry index is also shown as a black cross for each age group. For WTA, temporal, parietal, frontal, occipital lobes correspond to clusters 1, 2, 3, and 4 respectively, and limbic areas to cluster 5. Cluster colors correspond to those used in qualitative Figures (1,4 and 5)

In the WTA approach (Figure 7b), except the two dominant frontal and parietal clusters that are generally symmetrical. The symmetry of other clusters is very low (around 0.19 overall) compared to KNIT (0.58). Moreover, this method does not fully capture the increased symmetry through age.

##### Ipsilateral and contralateral connectivity symmetry

We did not observe any specific trend (Figure 8) of an increased nor decreased ispilateral nor contralateral connectivity as a function of age.

**Figure 8:**
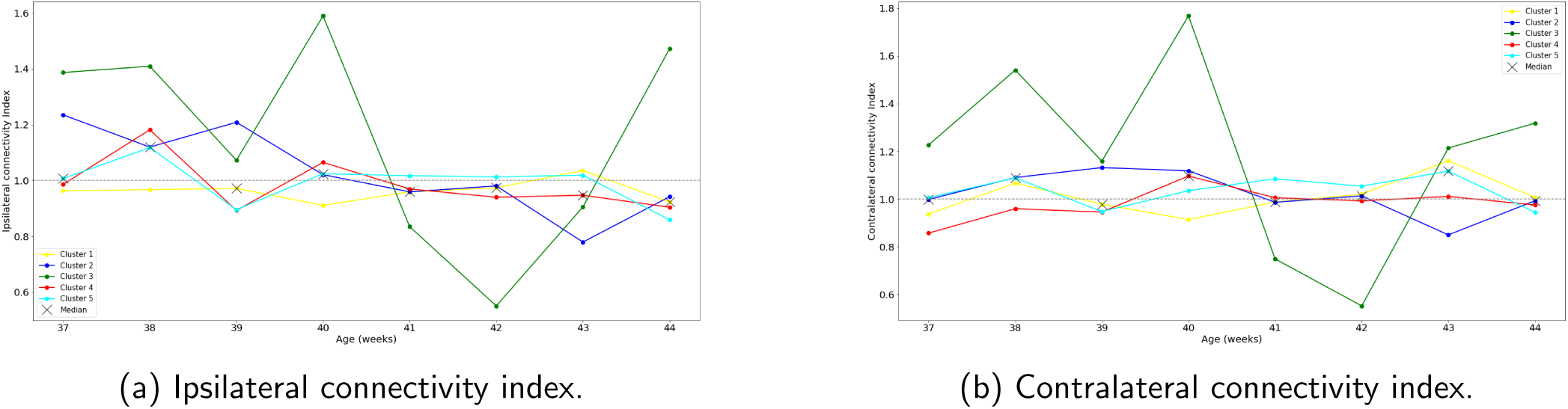
Ipsilateral and contralateral connectivity indices quantifying the symmetry and asymmetry respectively between thalamic connectivity to each hemisphere (as defined in Section 2.4).

##### Global thalamic connectivity

Global functional connectivity (Figure 9), i.e. independently of the clusters, shows a slow increase across age between 37 and 40 post-menstrual weeks, with a sharp increase at around 40 weeks, followed by a decrease of the connectivity to some areas at around 42 weeks. The areas that are highly connected to the thalamus are the parietal lobe, that hosts the somatosensory cortex, the superior temporal gyrus that contains the auditory cortex. This is line with the fetal development that involves more touch and auditory perceptions compared to other senses such as vision. The cingulate gyrus also shows high thalamic connectivity, along with the the frontal lobe that contains both the motor areas and the prefrontal cortex that is involved in planning and high order functions. The regions that are less connected to the thalamus contain the anterior temporal lobe, the amygdala and the hippocampus. The most significant surge appears to occur in the cerebellum (along with the parietal cortex). Although it is among the least connected regions during early developmental stages, a pronounced increase is observed at 41 weeks.

**Figure 9:**
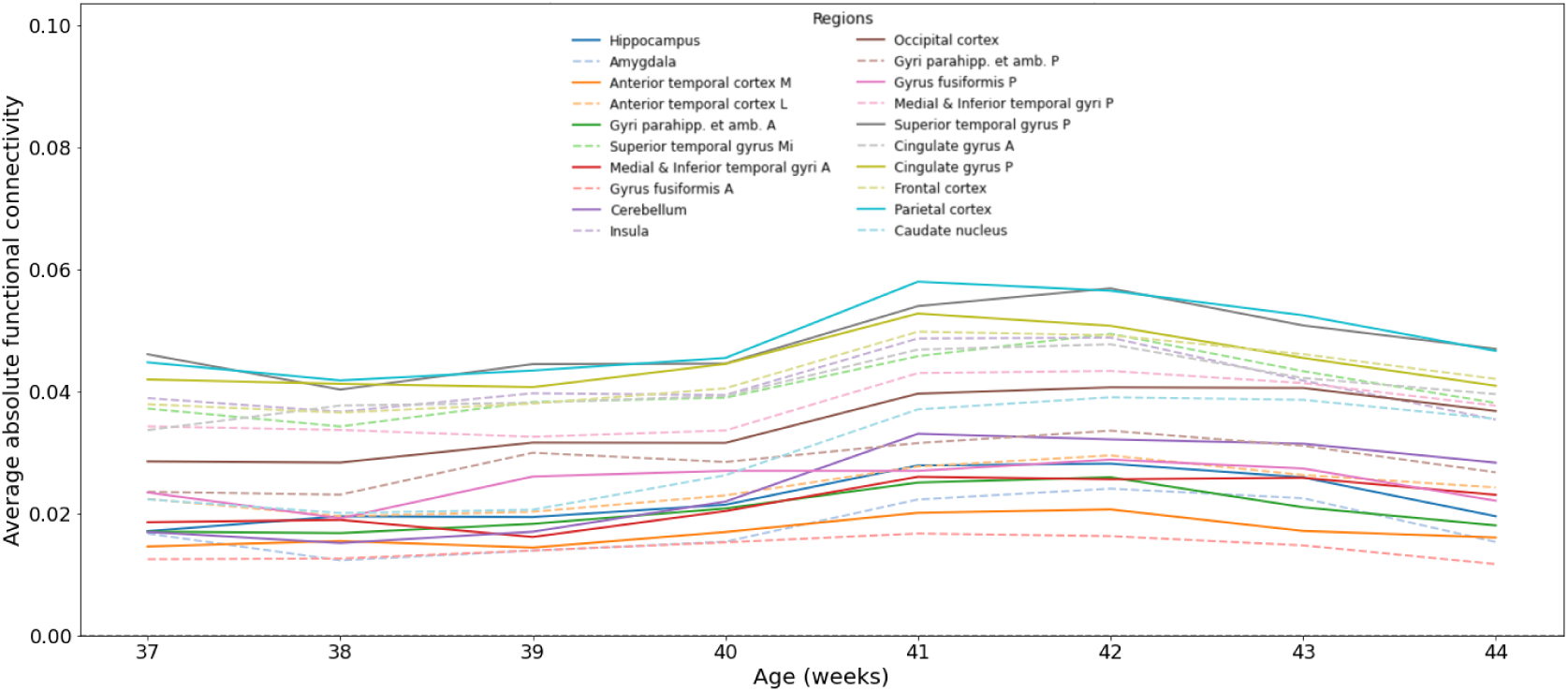
Global connectivity between the whole thalamus and the different ROIs (averaged bilaterally) across the age span.

##### ROI-ROI connectivity

We observe similar patterns in connectivity matrices of the different age groups with an increased connectivity for frontal, parietal and cingulate cortices to different brain areas and more prominently to their bilateral counterparts, as can be observed on the top left panel of Figure 10 for the 44 age group and in Supplementary Figure 17 for other age groups. However, whereas the superior temporal gyrus is a highly connected area to the thalamus (in the top 3 regions), its connectivity to other brain areas is not significant, as it only ranks on average (across age groups) in the 9th position. This shows that certain connectivity patterns are not necessarily driven by the whole brain connectivity but are specific to the thalamus.

**Figure 10:**
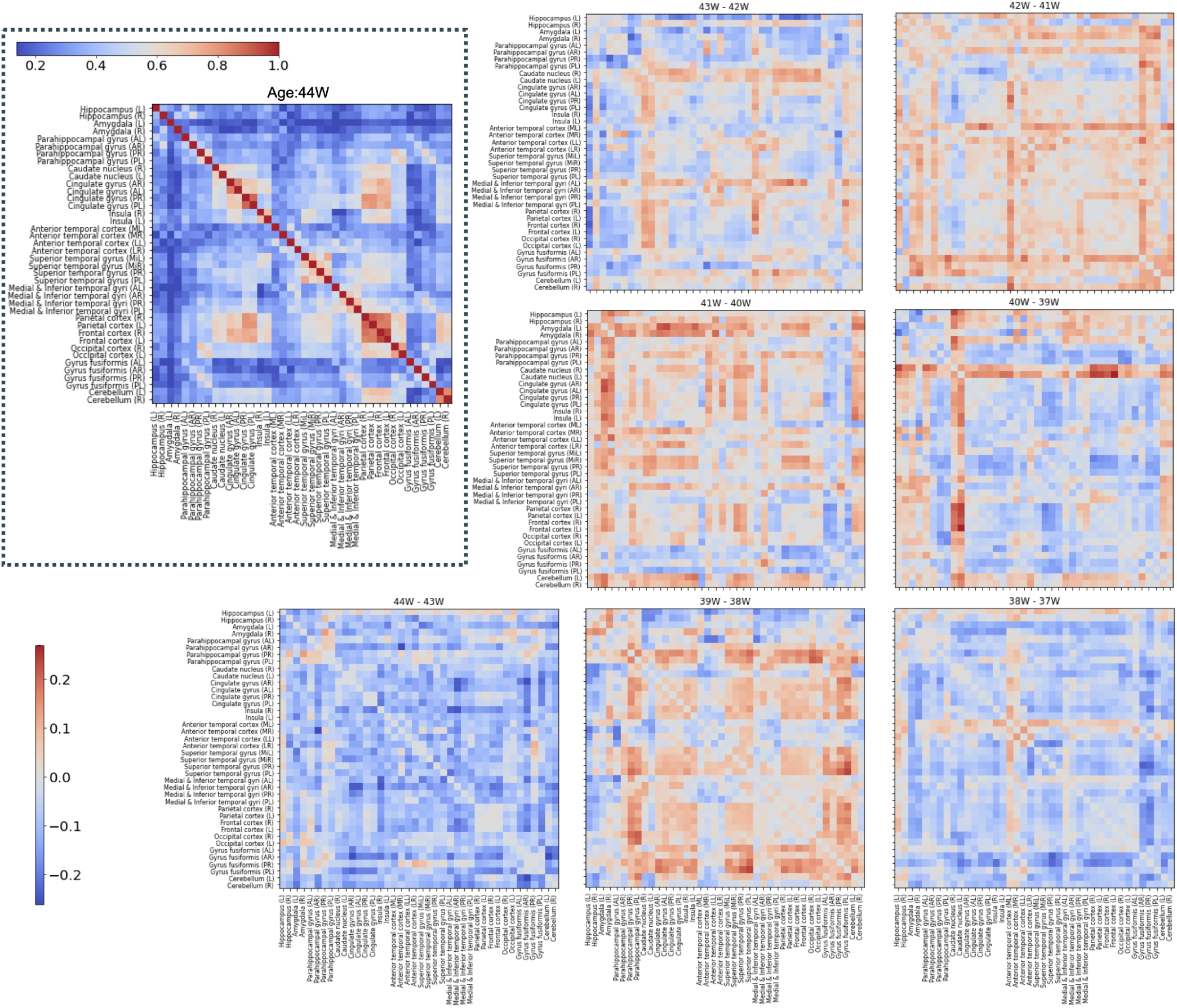
Top left panel: Functional connectivity matrix between the 40 ROIs. Other matrices show the differences between consecutive functional connectivity matrices from 44 to 37 weeks of age with a one week increment.

**Figure 11:**
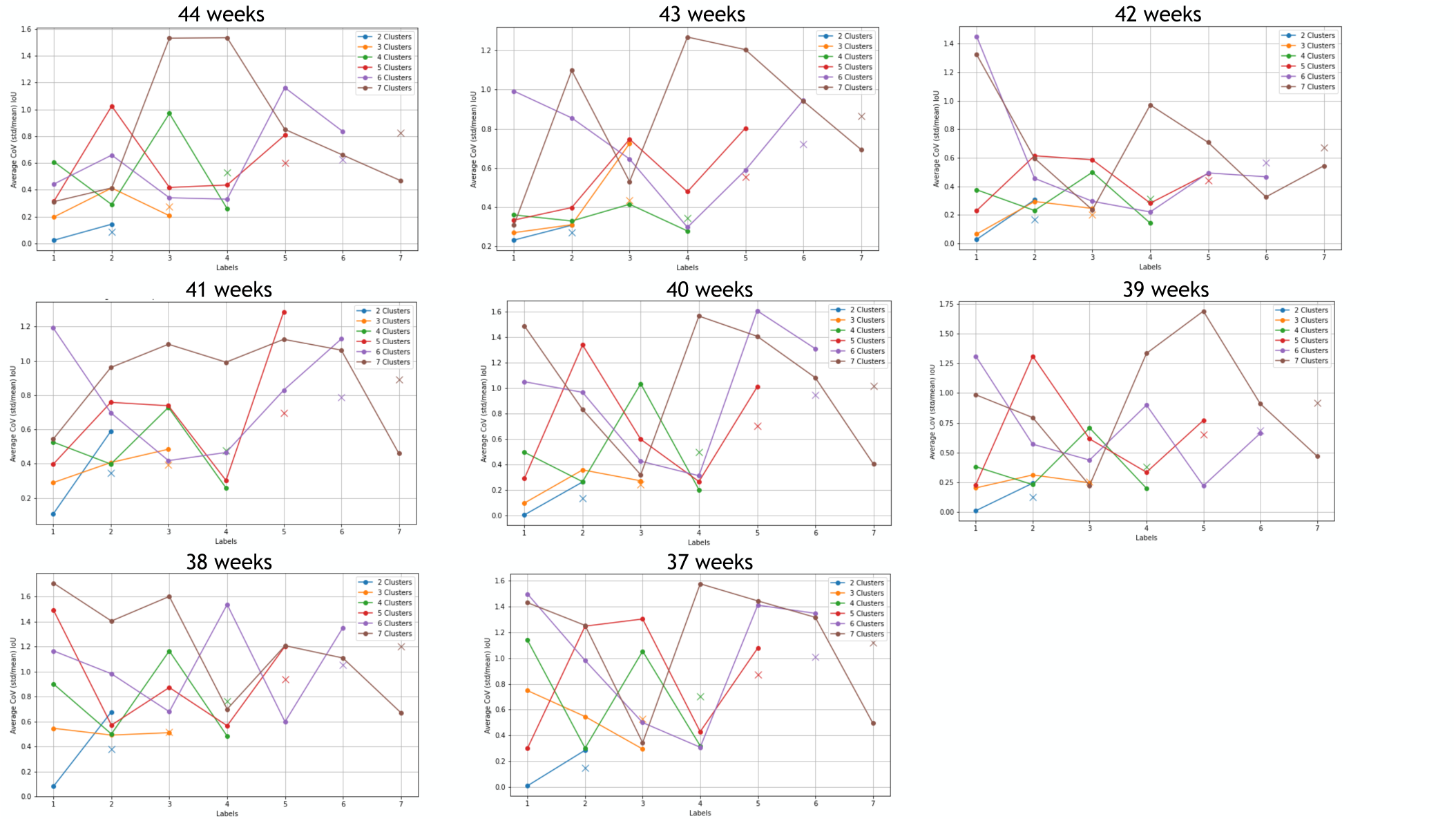
Variability through timepoints bootstraps for all age groups (N=25).

In terms of variation of FC matrices across the age span, we observe a general increase in connectivity that starts smoothly from the 39th week and starts decreasing from the 43rd week. This is similar to some extent to the thalamic connectivity to the different ROIs (Figure 9) and therefore part of this trend can be driven by this whole brain connectivity pattern (although it is more abrupt in the ROI-ROI case as can be seen in the supplementary Figure 18).

Specifically, we can notice for example that from 42 to 43 weeks we start to have an increase in connectivity in many regions (except limbic areas), especially the parietal, frontal, occipital, and cingulate regions to the caudate nucleus, which can explain the increase in dorsal cluster 4. volume The same cluster decreases in size from 41 to 42 weeks and this can also be explained by the decrease (blue color) in this connectivity pattern in the ROI-ROI matrix (42W-41W in Figure 10) compared to an increase in almost all other pairs (red color).

## 4 Discussion

In this study, we have shown how a fine-grained clustering framework of the thalamus (KNIT), using extensive connectivity information from a rich research dedicated dataset (Edwards et al., 2022), with 730 subjects acquired with 2300 temporal volumes, gives rise to spatially coherent and topologically symmetrical thalamic units across the age span of the developing neonates. In the following section we discuss the results obtained from this method with help from the current literature as well as the possible future avenues.

### 4.1 Functional parcellation by KNIT

By applying KNIT to connectivity profiles derived from 40 cortical and subcortical regions, we obtained 4 thalamic nuclei corresponding spatially and functionally to the pulvinar, ventrolateral, medial and dorsal thalamus. Here, KNIT uses extensive connectivity profiles rather than one-to-one connections (i.e. WTA approach) to inform the clustering algorithm of for the parcellation. Our results provides evidence that additional information may be deducted from these connectivity profile.

For instance, in terms of positive connectivity, the cluster 5 connects to parietal and superior temporal gyrus, additionally it connects to frontal and insular cortex. In that sense, it has a similar connectivity profile to cluster 2, except that cluster 2 has a lower connectivity to limbic areas compared cluster 5. This information helps the clustering algorithm to segregate cluster 5 from cluster 2 despite their physical proximity and the fact that they both have their strongest connections to the same regions.

Neurobiologically, the combination of strong and weak connections that a connectivity profile may provide may be indicative of functionality of the circuit that nucleus involves. In the case of cluster 2 and 5, the connectivity profile of cluster 2 (frontal, insula and anterior cingulate cortex) with weakest connectivity to the limbic system, may be indicative of the formation of the functional unit with the salience network and therefore the prefrontal cortex portion of the frontal lobe and the anterior part of the insula, in sync with anterior cingulate cortex and medial thalamus. Conversely, the connections of cluster 5, namely frontal and insula with relatively weaker (but not weaker than cluster 2, also in other age groups, i.e. Supplementary Figure 16) connections to the limbic system may be indicative of connectivity of the ventrolateral thalamus with the motor part of the frontal lobe and the posterior insula, with connections to limbic systems, helping in functions such as movement learning.

It is in that sense, we propose informing clustering algorithms of connectivity profiles and not one-to-one connections (i.e. WTA approach) in segmenting the functional units where complex connections such as thalamus are expected.

### 4.2 Early development of functional thalamic parcels

Delineated thalamic nuclei at 44 post-menstrual weeks showed a general increased connectivity to the superior temporal gyrus (auditory cortex), to the parietal lobe (somatosensory cortex) and to the frontal lobe (motor and executive areas). This *all* − *to* − *many* pattern shows a reduced specialization of the thalamus at a neonatal age as also demonstrated by Alcauter et al., 2014 where the sensorimotor cluster covers a large part of the thalamus including both the auditory and the visual areas. Similarly, the primary sensorimotor areas (Toulmin et al., 2015) and the motor area (Toulmin et al., 2021) were found to be significantly connected to the whole thalamus. However, a level of specificity was observed in our analysis where the pulvinar cluster was clearly delineated (as in Ferradal et al., 2019; Toulmin et al., 2015).

A ventrolateral cluster, that was shrinking with age, covering motor areas was also shown by our clustering algorithm. In Alcauter et al., 2014; Y. Cai et al., 2017, the sensorimotor cluster that was found significantly connected to the thalamus covers both our pulvinar and the ventrolateral clusters’ areas. This might indicate that our approach with a more fine-grained segmentation strategy and significantly higher number of subjects helps to delineate more specifically the thalamic nuclei. A medial cluster was also found in our results to be highly connected to the salience network. This network was reported to be one the two most significantly connected to the thalamus for neonates (Alcauter et al., 2014; Y. Cai et al., 2017) and its medial location is in line with our results.

Finally, the dorsal thalamus (cluster 4) that increased in volume and strength over age, showed a marked connectivity with the frontal and parietal cortices, possibly reflecting early development of the frontoparietal network (FPN). This circuit likely supports foundation of cognitive processes such as attention and working memory arising in early development. In this sense, dorsal thalamus may have a critical role relaying higher-order neural functions during the neonatal phase.

We also looked at the robustness of our clusters across age, two measures of uncertainty and symmetry were used with uncertainty decreasing through age and symmetry increasing. Briefly, our results indicate that from 37 to 44 weeks, spatial organization of the thalamus becomes more symmetric and functional clusters become more reproducible. Therefore, these results could hint towards an increase of functional specialization of the thalamus across the age span. We have also explored the ipsilateral and contralateral thalamic connectivity to other brain regions, but no specific trend has been found. However, this might be interesting to assess across diseases originating perinatally as this type of connectivity has been shown to be disrupted in schizophrenia for example (Liu et al., 2022).

We have also explored the global thalamic connectivity to other cortical and subcortical areas and found a transient increase at around 40 weeks of age. This can be in line with a recent longitudinal study (L. Ji et al., 2024) spanning both fetal and neonatal age (25 to 55 weeks) that found a burst in general connectivity at birth (i.e. 40 weeks).

Finally, examining the overall brain connectivity revealed that thalamic connectivity patterns are driven by the dynamics of the entire brain, particularly those involving the frontal, parietal, and cingulate regions. In contrast, the connectivity of the superior temporal gyrus appears to be specific to the thalamus.

### 4.3 Future perspective

The transition from intra-auterine to extra-auterine life involves abrupt environmental changes that imposed evolutionary adaptation strategies to the physiology of both the brain and body (Hillman et al., 2012). In future work, we would like to extend our study to the fetal period (Taymourtash et al., 2022, 2023) using the new release of the fetal dHCP dataset (Karolis et al., 2024).

Moreover, how structural parcellations of the thalamus with diffusion MRI and tractography relate to the functional parcellations with rs-fMRI and functional connectivity can be a promising area of work (as explored by Ferradal et al., 2019). In fact, it is hypothesized (Ferradal et al., 2019) that the agreement between structural and functional connectivity maps will show a higher overlap in sensory systems compared to higher-order cognitive regions. Exploring this hypothesis across age or disease can be highly interesting. Several studies have already explored the thalamic connectivity with diffusion MRI in the perinatal period by characterizing development or by comparing full-term to pre-term or to congenital heart disease newborns (Jaimes et al., 2018; Jakab et al., 2020; H. Ji, Payette, et al., 2024; H. Ji, Wu, et al., 2024; Oldham et al., 2024; Zheng et al., 2023). For instance, Zheng et al., 2023 have found using 175 subjects of the dHCP (same dataset as used in our work) spanning 32-44 post-menstrual weeks, with a WTA approach on the number of white matter streamlines to 5 cortical regions, an overall symmetrical parcellation with an overall highly increasing frontal and somatosensory nuclei across the age span, as also demonstrated by our study. Another way of quantifying structure and function interactions can be through the structural-decoupling index (Preti & Van De Ville, 2019) using graph signal processing, as performed in adult connectomes. Finally, and if specific absorption rate (SAR) challenges (Bridgen et al., 2024; Malik et al., 2021) were addressed, imaging in high field (7T MRI) that drastically increases the signal-to-noise ratio (SNR) can exhibit more robust clusters as has been performed for adult brains (Jorge et al., 2020; Martinez et al., 2024). In fact, a higher resolution can help us delineate more thalamic subnuclei, especially for older neonates, as shown by our bootsrapping and uncertainty metrics.

The neonatal data acquired with EPI suffer from low SNR and from motion residual artefacts (despite correction, as has been shown in Kebiri et al., 2024). Nonetheless, the findings of our study provide valuable insights into the developmental connectivity of the thalamus. By leveraging a large cohort, a high dimensional feature space in clustering, along with bootstrapping and uncertainty techniques, we addressed the challenges associated with neonatal data quality and captured meaningful developmental patterns.

These findings lay a foundation for future investigations into how thalamic functional organization evolves across critical periods of brain maturation and in the context of perinatal disorders. Continued exploration in that direction has the potential to advance our understanding of the role of the thalamus in early-life brain organization and its impact on neurodevelopmental outcomes.

## Ethics Statement

This study uses pre-approved dHCP data by UK Health Research Authority (REC reference: 14/LO/1169), requiring no additional ethical approval.

## Acknowledgements

We gratefully acknowledge access to the facilities and expertise of the CIBM Center for Biomedical Imaging (Centre d’Imagerie BioMédicale), a Swiss research center of excellence founded and supported by Lausanne University Hospital (CHUV), University of Lausanne (UNIL), EÉcole Polytechnique Fédérale de Lausanne (EPFL), University of Geneva (UNIGE), Geneva University Hospitals (HUG), and the Leenaards and Jeantet Foundations.

This research was funded by grants from the Swiss National Science Foundation (grants 182602, 215641 and PZ00P2 185909). It was also supported by CSEM – Swiss Center for Electronics and Microtechnology, by the Translational Imaging Center (TIC) of the Swiss Institute for Translational and Entrepreneurial Medicine (SITEM).

The authors are grateful to the TANGO consortium members (https://thalamicsegmentation.github.io/), Dr. Vinod Kumar and Prof. Manoj Saranathan, for the exceptional opportunities for scientific discussions which contributed towards this work.

## A Appendix

**Figure 12:**
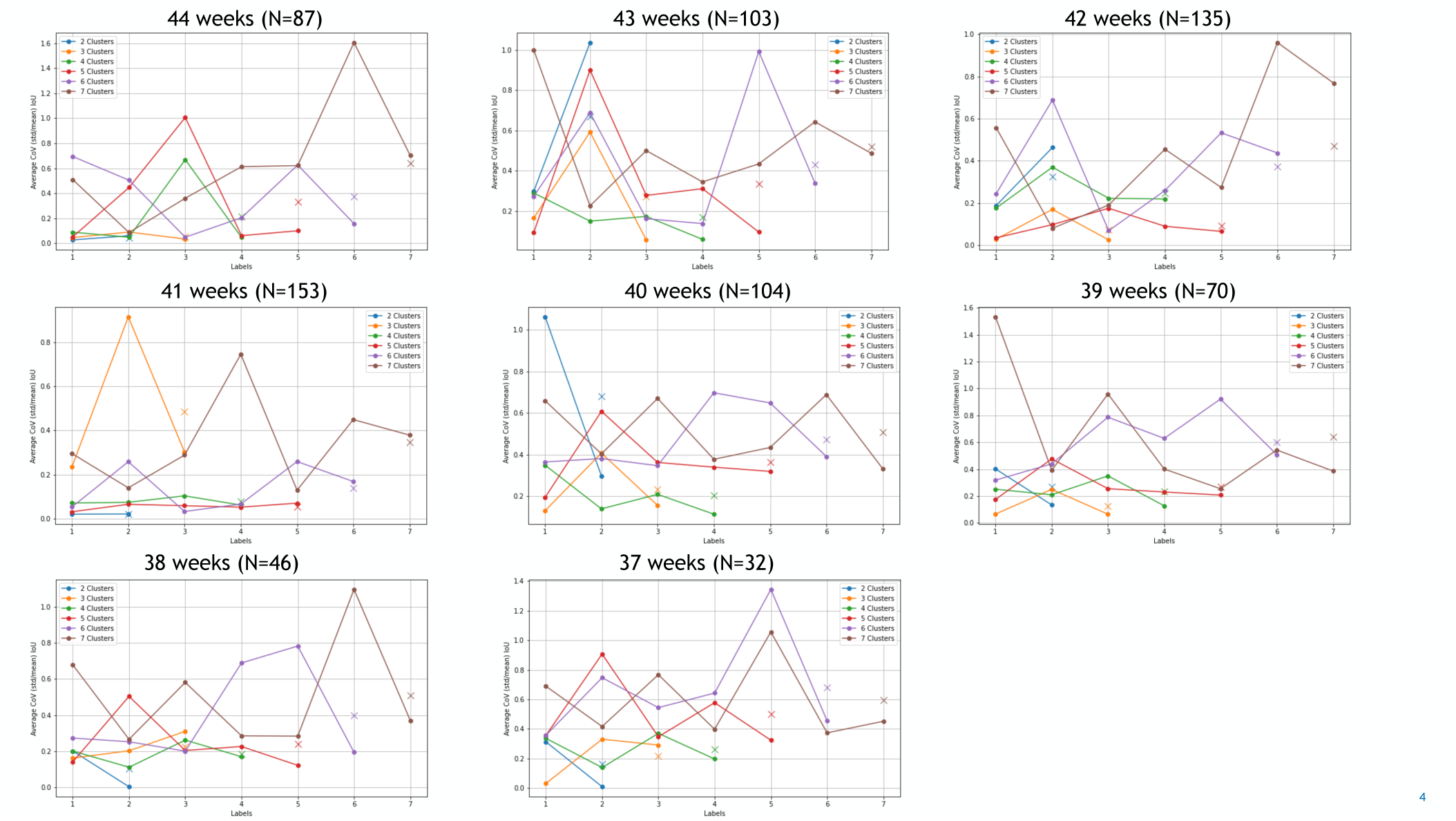
Variability through subject bootstraps for all age groups. The number of subjects per age group is shown on the top.

**Table 2:** Mapping of the 40 seed ROIs and their respective abbreviations used throughout the manuscript. The last four rows correspond to the thalamic labels that were merged to define thalamic voxels.

| ROI Label Name in dHCP | ROI Abbreviation |
| --- | --- |
| Hippocampus left | Hippocampus (L) |
| Hippocampus right | Hippocampus (R) |
| Amygdala left | Amygdala (L) |
| Amygdala right | Amygdala (R) |
| Anterior temporal lobe, medial part left GM | Anterior temporal cortex (ML) |
| Anterior temporal lobe, medial part right GM | Anterior temporal cortex (MR) |
| Anterior temporal lobe, lateral part left GM | Anterior temporal cortex (LL) |
| Anterior temporal lobe, lateral part right GM | Anterior temporal cortex (LR) |
| Gyri parahippocampalis et ambiens anterior part left GM | Parahippocampal gyrus (AL) |
| Gyri parahippocampalis et ambiens anterior part right GM | Parahippocampal gyrus (AR) |
| Superior temporal gyrus, middle part left GM | Superior temporal gyrus (MiL) |
| Superior temporal gyrus, middle part right GM | Superior temporal gyrus (MiR) |
| Medial and inferior temporal gyri anterior part left GM | Medial & Inferior temporal gyri (AL) |
| Medial and inferior temporal gyri anterior part right GM | Medial & Inferior temporal gyri (AR) |
| Lateral occipitotemporal gyrus, gyrus fusiformis anterior part left GM | Gyrus fusiformis (AL) |
| Lateral occipitotemporal gyrus, gyrus fusiformis anterior part right GM | Gyrus fusiformis (AR) |
| Cerebellum left | Cerebellum (L) |
| Cerebellum right | Cerebellum (R) |
| Insula right GM | Insula (R) |
| Insula left GM | Insula (L) |
| Occipital lobe right GM | Occipital cortex (R) |
| Occipital lobe left GM | Occipital cortex (L) |
| Gyri parahippocampalis et ambiens posterior part right GM | Parahippocampal gyrus (PR) |
| Gyri parahippocampalis et ambiens posterior part left GM | Parahippocampal gyrus (PL) |
| Lateral occipitotemporal gyrus, gyrus fusiformis posterior part right GM | Gyrus fusiformis (PR) |
| Lateral occipitotemporal gyrus, gyrus fusiformis posterior part left GM | Gyrus fusiformis (PL) |
| Medial and inferior temporal gyri posterior part right GM | Medial & Inferior temporal gyri (PR) |
| Medial and inferior temporal gyri posterior part left GM | Medial & Inferior temporal gyri (PL) |
| Superior temporal gyrus, posterior part right GM | Superior temporal gyrus (PR) |
| Superior temporal gyrus, posterior part left GM | Superior temporal gyrus (PL) |
| Cingulate gyrus, anterior part right GM | Cingulate gyrus (AR) |
| Cingulate gyrus, anterior part left GM | Cingulate gyrus (AL) |
| Cingulate gyrus, posterior part right GM | Cingulate gyrus (PR) |
| Cingulate gyrus, posterior part left GM | Cingulate gyrus (PL) |
| Frontal lobe right GM | Frontal cortex (R) |
| Frontal lobe left GM | Frontal cortex (L) |
| Parietal lobe right GM | Parietal cortex (R) |
| Parietal lobe left GM | Parietal cortex (L) |
| Caudate nucleus right | Caudate nucleus (R) |
| Caudate nucleus left | Caudate nucleus (L) |
| Thalamus right, high intensity part in T2 | Thalamus (R) |
| Thalamus right, low intensity part in T2 | Thalamus (R) |
| Thalamus left, high intensity part in T2 | Thalamus (L) |
| Thalamus left, low intensity part in T2 | Thalamus (L) |

**Figure 13:**
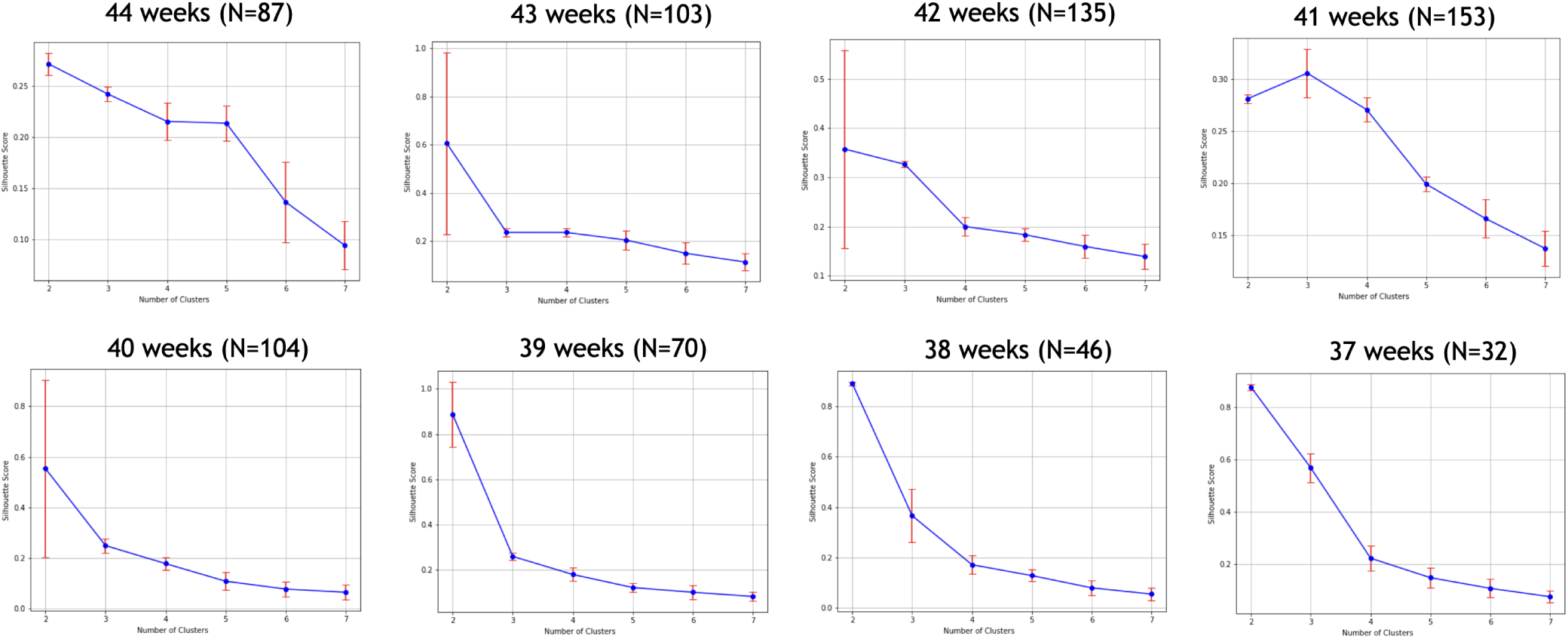
Silhouette score for the different cluster numbers (*k*) across all age groups. The number of subjects per age group is shown on the top.

**Figure 14:**
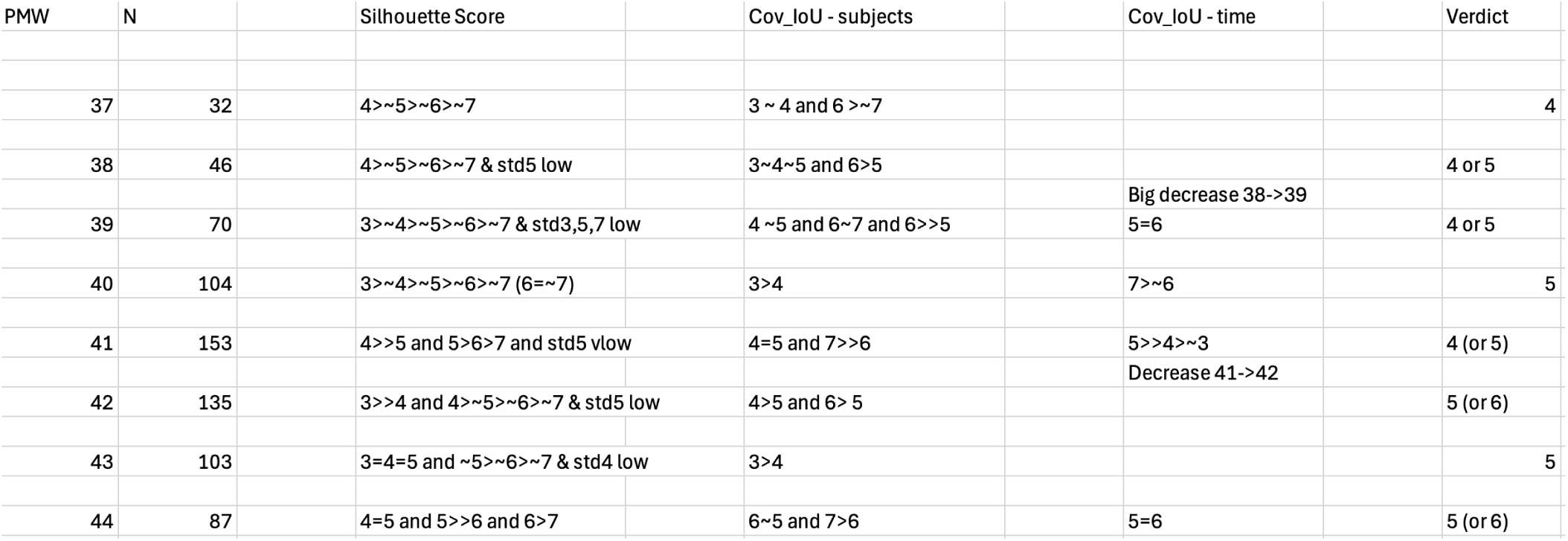
Heuristics on the choice of the number of clusters *k* for k-means in KNIT.

**Figure 15:**
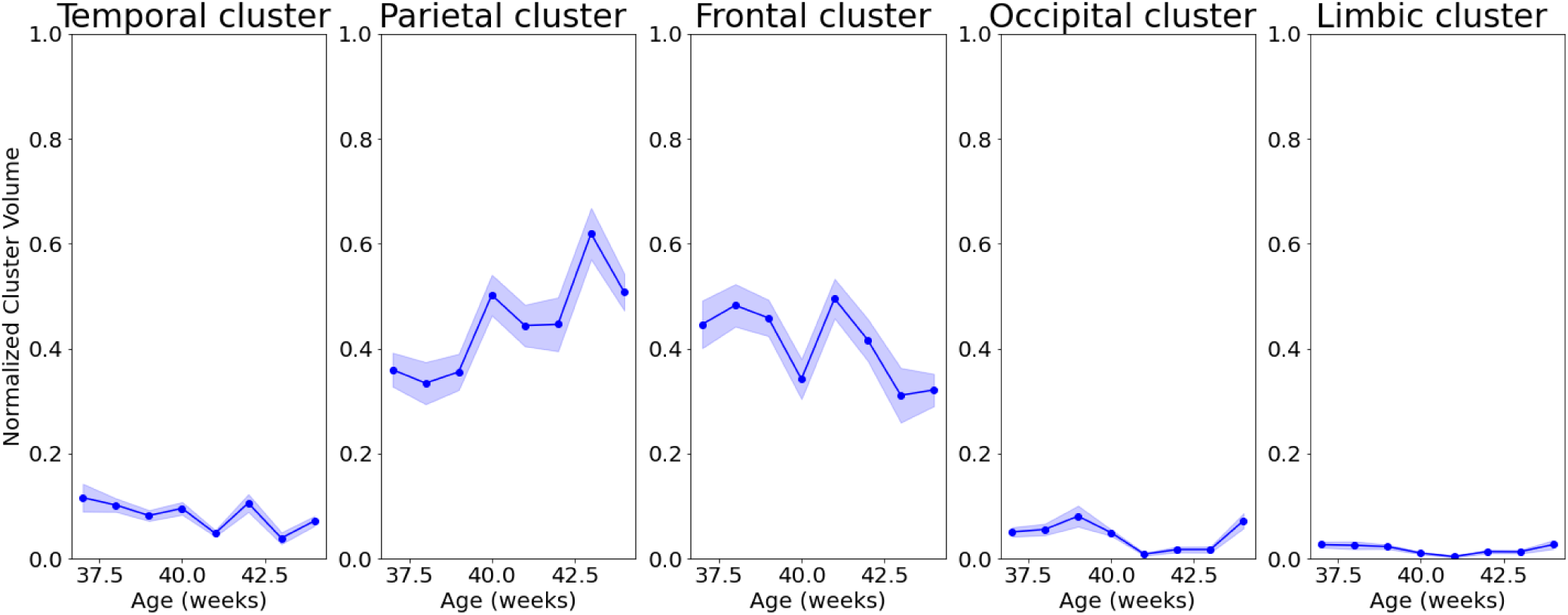
Developmental evolution of the normalized volumes (across whole thalamic volume) for each cluster, for the WTA approach. Mean and standard deviations across subject bootstraps are displayed.

**Figure 16:**
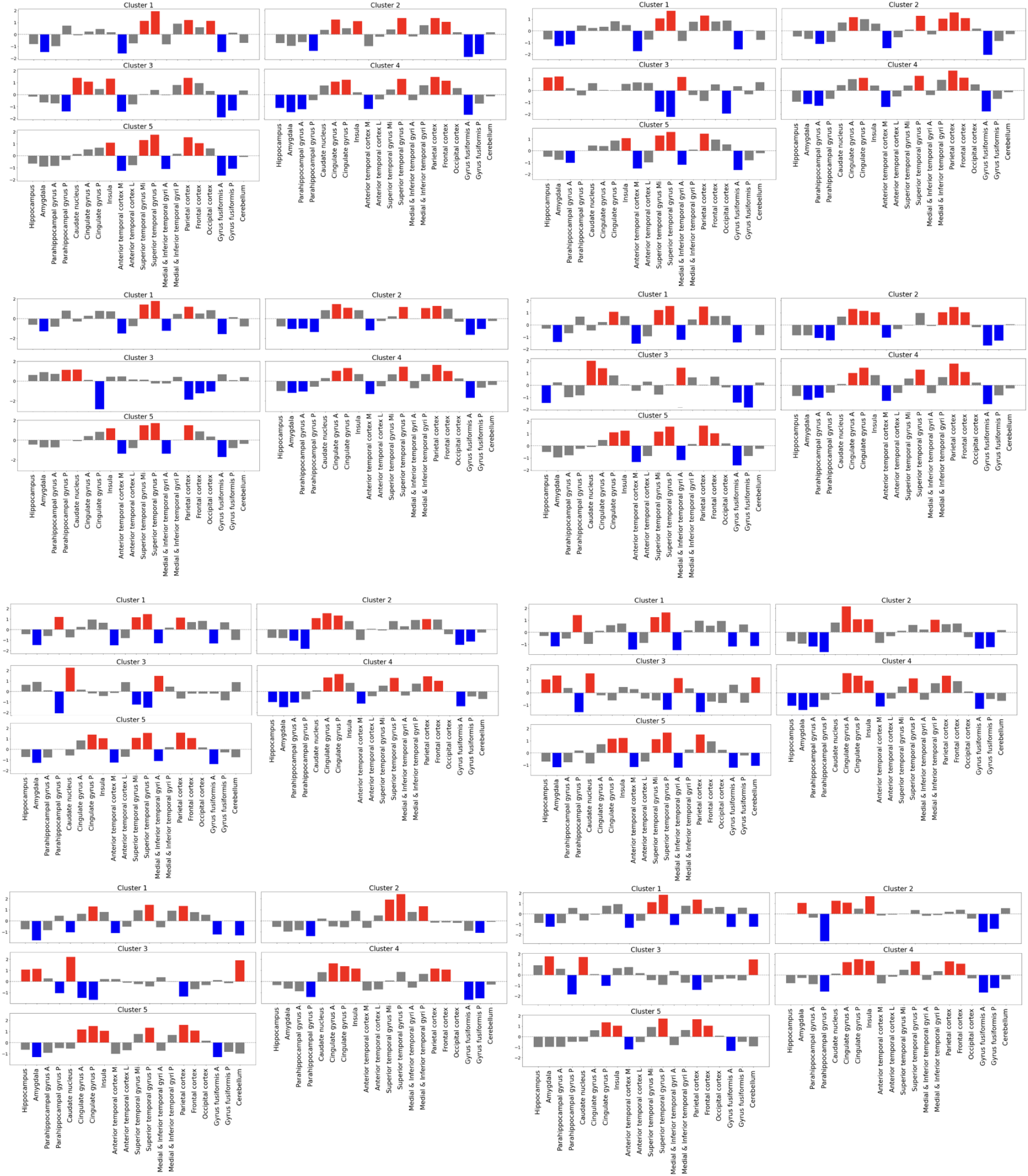
Z-score of the median FC, between each cluster ID and the 40 bilateral brain regions that were merged between the right and the left hemispheres, for all age groups. From 44 weeks on the top left to the 37 weeks age group on bottom right.

**Figure 17:**
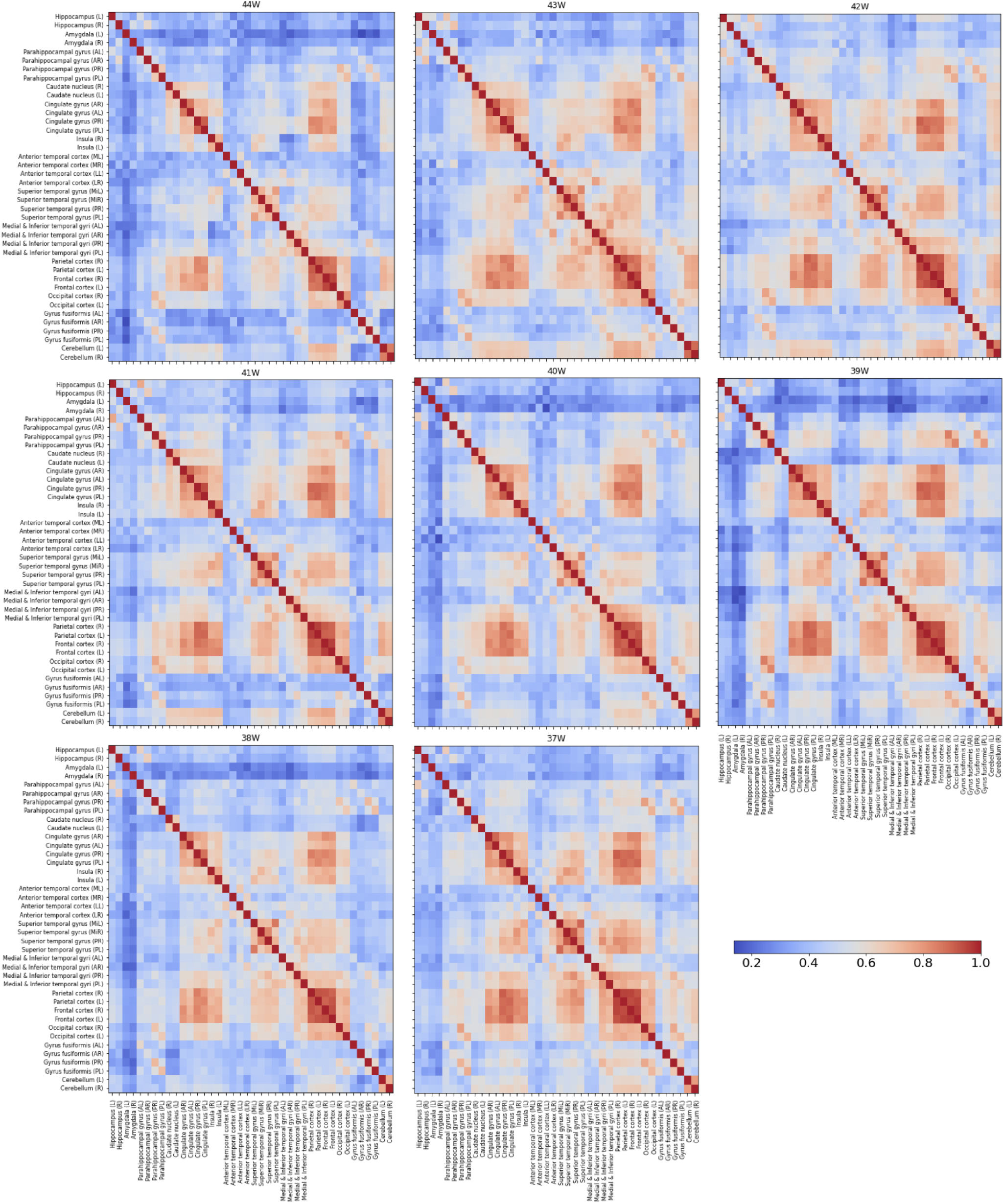
Functional connectivity matrices between the 40 ROIs of the different age groups.

**Figure 18:**
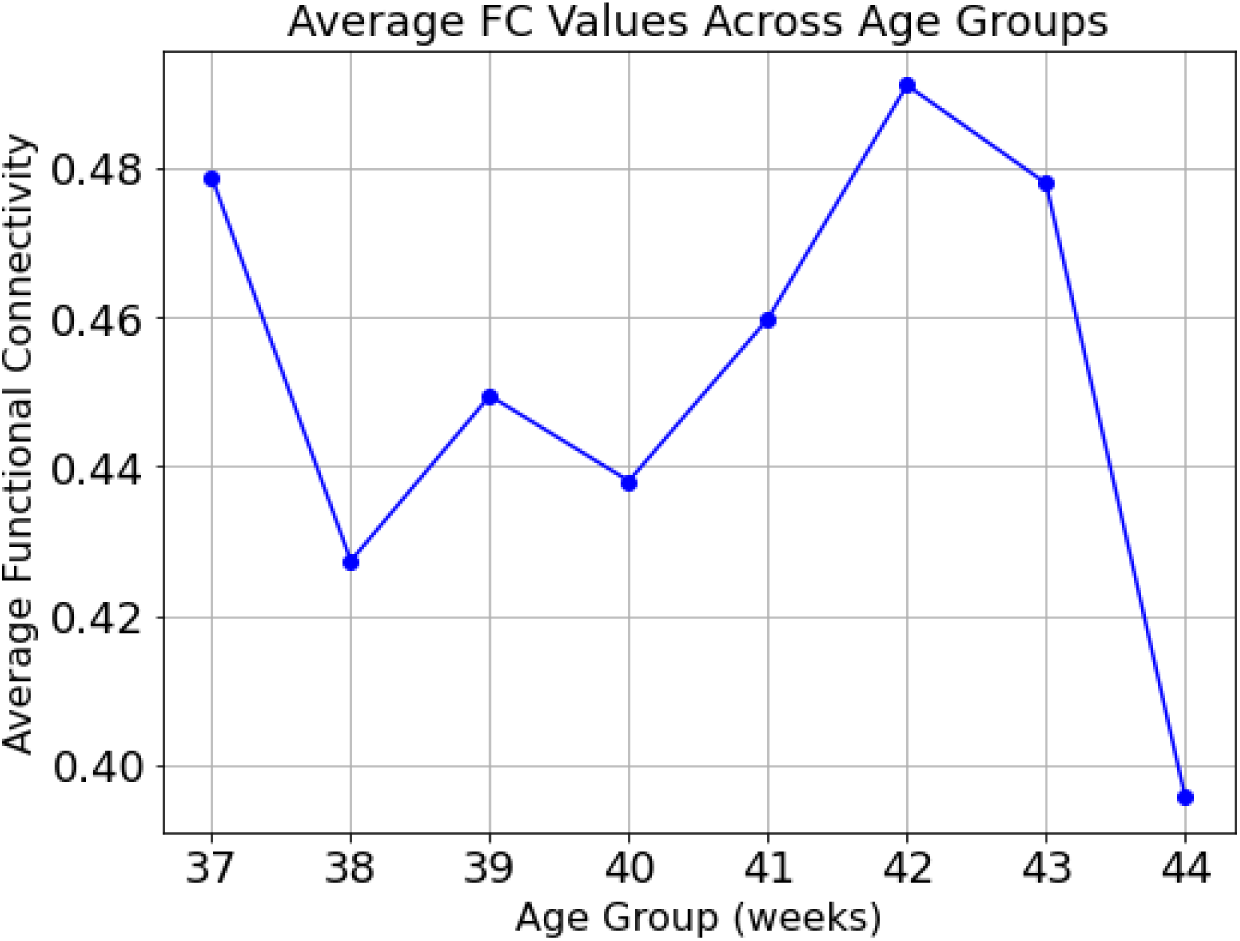
Average ROI-ROI functional connectivity evolution across different age groups.

## Footnotes

1 https://www.developingconnectome.org/data-release/third-data-release/

